# Cytotoxic CD4 Development Requires CD4 Effectors to Concurrently Recognize Local Antigen and Encounter Infection-Induced IL-15

**DOI:** 10.1101/2020.09.03.281998

**Authors:** Priyadharshini Devarajan, Allen M. Vong, Catherine H. Castonguay, Noah J. Silverstein, Olivia Kugler-Umana, Bianca L. Bautista, Karen A. Kelly, Jeremy Luban, Susan L. Swain

## Abstract

Cytotoxic CD4 T cells (ThCTL) are tissue-resident effectors that enhance viral clearance by MHC-II-restricted cytotoxicity of infected cells. Using a model of influenza A virus (IAV) infection, we identify key factors that drive CD4 effectors to differentiate into lung-resident ThCTL. We find that, to become ThCTL, CD4 effectors must again recognize cognate antigen on antigen presenting cells (APC) within the lung. Different APC subsets can drive this transition, including dendritic cells, B cells, and to a lesser extent non-hematopoietic MHC-II^+^ APC. CD28 co-stimulation is not required and can reduce ThCTL development. In contrast, T follicular helper cells (T_FH_) that are another specialized CD4 effector subset, require CD28 during this time. Optimal ThCTL generation also requires ongoing infection in the effector phase, that acts independently of antigen presentation. The mechanism involves production of Type I IFN, that induces IL-15 which acts to support further differentiation of CD4 effectors to ThCTL. The multiple spatial, temporal and cellular requirements for ThCTL generation from CD4 effectors described here would be expected to prevent cytotoxic CD4 responses in the lung after pathogen has already been cleared, while ensuring the development of potent lung-restricted ThCTL effectors when pathogen persists.

## INTRODUCTION

Among T cells, CD8 cells are thought of as the cytotoxic lineage. However, in the past decade, many studies have shown that cytotoxic CD4 T cells (ThCTL) can protect against intracellular pathogens and tumors but also propagate human diseases (Cenerenti et al., 2022; Oh and Fong, 2021). In the last year, their role has been highlighted in two human diseases, emphasizing the need to understand factors regulating their development. In one, long-term remission in cancer patients was linked to cytotoxic CD4 T cells induced by CAR-T therapy (Melenhorst et al., 2022). In the other, increased cytotoxic CD4 T cells in the lung and cytotoxic CD4 signatures in blood were found to correlate with worse outcomes in severe COVID-19 patients (Kaneko et al., 2022; Meckiff et al., 2020).

Various mouse models of infection have been used to study ThCTL in the infected tissue and their contributions to the anti-viral immune response including mousepox, CMV (LCMV and MCMV) and influenza infection (Knudson et al., 2021; Marshall et al., 2016; Wehrens et al., 2018). ThCTL can play an important role in controlling primary infections using a perforin dependent mechanism to kill infected cells and can synergize with B cell responses to enhance immunity (Brown et al., 2006; Brown et al., 2012; Fang et al., 2012). This is especially important when viruses downregulate MHC-I as an immune evasion mechanism (Marshall and Swain, 2011; Soghoian and Streeck, 2010). However, overt ThCTL responses may also contribute to immunopathology as indicated by recent COVID-19 studies (Kaneko et al., 2022; Meckiff et al., 2020). Thus, it is crucial to define what signals drive ThCTL generation during infections so as to learn how to enhance them with vaccines or block them when they are harmful.

A mouse model of influenza infection allows us to study aspects of a successful T cell response because primary influenza infection is cleared by T cells with antibody responses developing only thereafter (Swain et al., 2006). Acute influenza infection induces lung-resident ThCTL beginning at 6-7 days post infection (dpi), identified by surface expression of NKG2C/E (Marshall et al., 2016). Only NKG2C/E-expressing CD4 effectors in the infected lung mediate perforin-dependent cytotoxicity specific for MHC-II-expressing target cells (Marshall et al., 2016). Consistent with their restricted location, ThCTL express a tissue-resident phenotype and are not present in uninfected peripheral tissues, such as the spleen and draining lymph node (DLN) during influenza infection (Marshall et al., 2016). Most previous studies used granzyme B (GzmB) as the signature marker for cytotoxic CD4 T cells, however the majority of CD4 effectors in the inflamed tissue express GzmB, even though they are not all functionally cytotoxic (Marshall et al., 2016). GzmB is also associated with processing of proinflammatory cytokines and has various additional non-cytotoxic functions (Richardon et al., 2022). Thus, we can reliably use NKG2C/E to identify ThCTL, to unambiguously track the generation of ThCTL effectors and determine what signals are needed for their development.

The formation of cytotoxic CD4 T cells in response to infection seems to be a two-step process: (1) the generation of Th1 CD4 effectors and (2) the further differentiation of some of these into cytotoxic CD4 effectors (Knudson et al., 2021; Krueger et al., 2021). However, the mechanisms and signals needed to drive the transition of CD4 effectors into ThCTL, and where they are needed are unclear. Here we use a sequential adoptive transfer model to pinpoint the roles of antigen (Ag) recognition, costimulatory signals, and other signals from infection at this 2^nd^ step, in the development of CD4 effectors generated during influenza infection into ThCTL in the lung. We find that CD4 effectors at 6 dpi need to recognize cognate antigen again, but this must occur in the lung to drive ThCTL development. Several APC subsets can effectively present Ag at this timepoint. This pathway does not require CD28 co-stimulation during antigen recognition, and it can be antagonistic. Finally, we find a requirement for infection-generated Type I IFN to induce IL-15, which synergizes with local Ag presentation to drive the development of full-fledged lung ThCTL. Thus, CD4 effectors develop into ThCTL only when multiple infection-generated signals are present, so that cytotoxic CD4 responses are induced only when infection is ongoing and not if virus has already been cleared. We discuss the relevance of these findings to vaccine and immunotherapeutic strategies for tumors and viral infections.

## RESULTS

### Cognate Ag recognition by CD4 effectors drives the generation of ThCTL phenotype and function in the lung

Following influenza infection, we find that all MHC-II-restricted functional cytotoxicity in the lung is mediated by NKG2C/E expressing CD4 effectors (Marshall et al., 2016). Thus, we can use surface staining for NKG2A/C/E as a signature marker to readily identify ThCTL and study the factors driving their generation, which previously had been challenging due to the lack of a reliable surface marker that clearly marked functionally cytotoxic CD4 T cells *in vivo*. During influenza infection, ThCTL develop only in the lung and after 6 days post-infection (dpi), peak at 8-10 dpi and contract thereafter (Marshall et al., 2016) (**Figure S1A**). To determine if Ag presentation after 6 dpi plays a role in ThCTL generation, we transferred naïve OT-II.Nur77^GFP^.Thy1.1^+^ CD4 T cells into wild-type (WT) hosts and infected them with PR8-OVA_II_. Nur77^GFP^ CD4 T cells transiently express Nur77 and thus GFP when they are stimulated by Ag recognition (Au-Yeung et al., 2014; Bautista et al., 2016; Moran et al., 2011). At 6 dpi, more of the Nur77^GFP+^ than the Nurr77^GFP-^ donor cells in the lung expressed NKG2A/C/E, and levels of Nurr77 were higher on NKG2A/C/E^+^ cells (**Figure S1B**), indicating that the ThCTL in the lung have recently recognized Ag.

ThCTL have been found to develop from already generated Th1 effectors (Knudson et al., 2021; Krueger et al., 2021). After influenza infection Th1 effectors traffic to the lung from the periphery, beginning at 6 dpi (Strutt et al., 2012). Therefore, we used a sequential transfer approach (Bautista et al., 2016) to determine if 6 dpi effectors, isolated from the spleens and DLN of infected host, require Ag recognition to become ThCTL (**Figure 1A**). We generated 6 dpi CD4 effectors *in vivo* by transfer of naïve OT-II (specific for an OVA_II_ epitope) Thy1.1 CD4 T cells, into 1^st^ hosts infected with PR8-OVA_II_ influenza A virus. We isolated the donor-derived *in vivo*-generated effectors from the spleen and DLN at 6 dpi. The 6 dpi effectors contain no ThCTL (Marshall et al., 2016), but we predict they include the CD4 effector precursors to lung ThCTL. These donor effectors were then transferred into 2^nd^ hosts that had been infected 6d previously (infection-matched) with or without Ag, to ensure that signals from infection other than cognate Ag, were reproduced as in a normal *in situ* response. This allows us to separate the 1^st^ step of CD4 effector generation from the 2^nd^ step during which ThCTL are generated from the CD4 effectors, in an *in vivo* system. We provided 2^nd^ hosts with either Ag and infection (PR8-OVA_II_ infection) or infection without Ag (PR8 infection) or neither (uninfected) (**Figure 1A**). We assessed donor ThCTL 2 days post transfer (dpt) in the lung, which corresponded to 8 dpi. A clear cohort of donor ThCTL developed only when Ag was present in the 2^nd^ hosts (PR8-OVA_II_), with few found when Ag was absent (PR8) (7-fold less) (**Figure 1B**). This Ag-dependence was more selective for ThCTL than the total CD4 effector population, as the total number of donor effectors in the lung was only reduced 1.7-fold without Ag (**Figure S1C**). Donor cell expression of other indicators of cytotoxicity, such as Granzyme B (GzmB) (**Figure 1C**) and the degranulation marker CD107a (**Figure 1D**), were also dependent on Ag presented in the 2^nd^ host. Expression of other ThCTL-associated markers (Marshall et al., 2016) by the donor lung effectors, PD1 and CXCR6 were also markedly Ag dependent, while two other markers, activated PSGL1 and decreased CXCR3 were not (**Figure S1D-G**).

**Figure 1.**
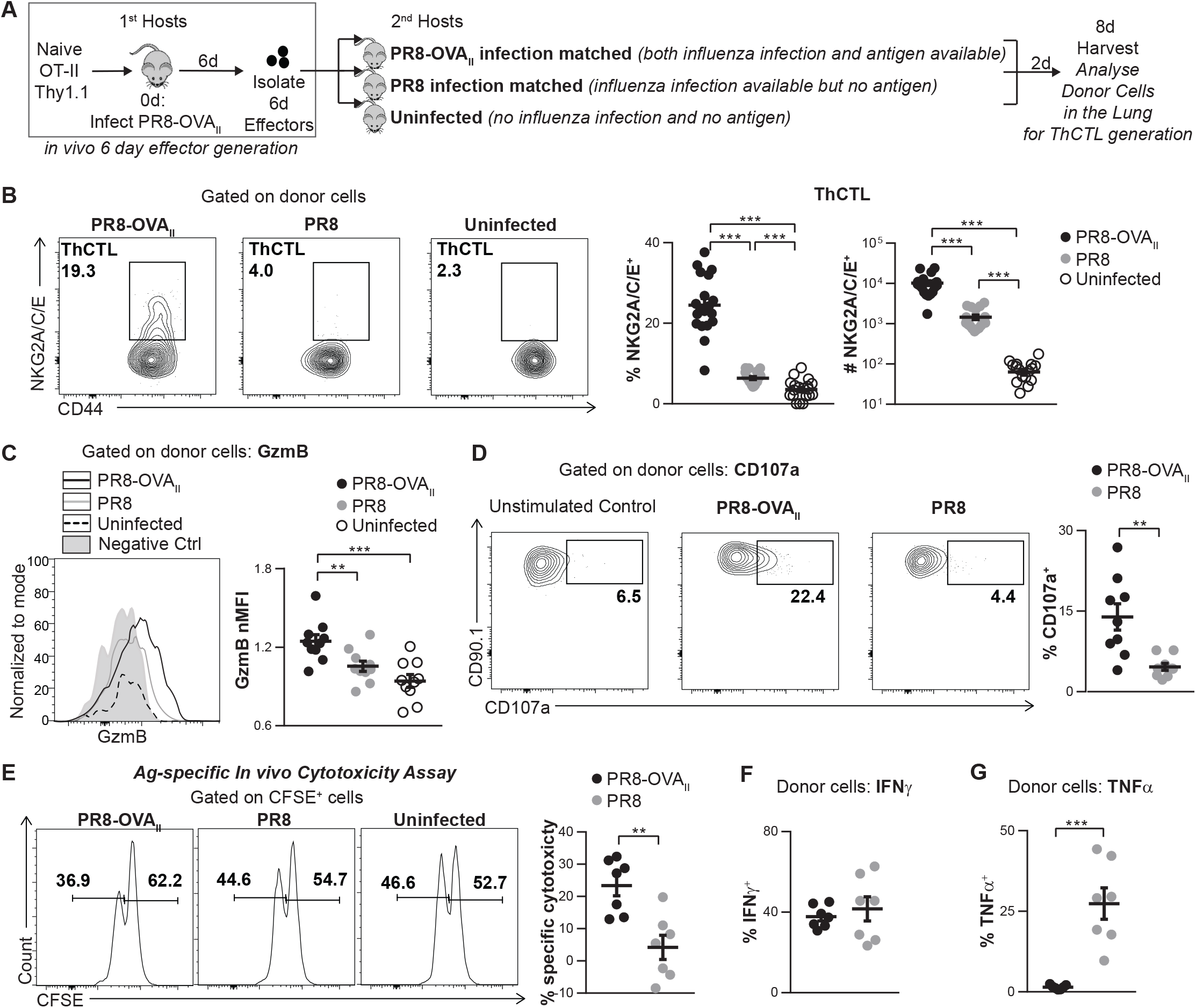
Cognate Ag during the effector checkpoint is required for lung ThCTL phenotype and function. **(A)** Experimental design for (B-D): Naïve OT-II.Thy1.1^+^ cells were transferred into PR8-OVA_II_ infected mice (1^st^ hosts). At 6 dpi, OT-II.Thy1.1^+^ effectors were isolated from 1^st^ hosts and transferred into following groups of 2^nd^ hosts: 6 dpi PR8-OVA_II_-infected, 6 dpi PR8-infected, or uninfected mice. Donor cells in the lung were analyzed 8 dpi. **(B)**Percentage and numbers of donor lung ThCTL (NKG2A/C/E^+^) (n=19 per group pooled, 4 independent experiments). **(C)** Representative histogram of donor lung cell GzmB expression (negative control: naïve CD4 from uninfected mice). Normalized MFI of donor lung cell GzmB expression (n=10 per group pooled, 2 independent experiments). **(D)** CD107a degranulation marker expression by donor lung cells (n=9 per group pooled, 2 independent experiments). **(E)** Experimental design: *In vivo* 6d OT-II.Thy1.1^+^ effectors were transferred into 6 dpi PR8-OVA_II_ or PR8 infection-matched TCRα/β^-/-^ mice. CFSE^lo^ target and bystander CFSE^hi^ bystander cells were transferred at 7d. Representative CFSE histograms shown. Percentage Ag specific cytotoxicity in each group is shown. **(F-G)** Experiment done as in (E). Percentage of donor lung cells expressing intracellular IFNγ (F) and TNFα (G) (E-G, n=7 per group pooled, 2 independent experiments). Statistical significance determined by two-tailed, unpaired Student’s t-test (* P<0.05, ** P<0.01, *** P<0.001). See also Fig. S1.

To assess *in vivo* cytotoxic function, we transferred *in vivo*-generated effectors into 2^nd^ hosts with or without Ag (as in Figure 1A) along with CFSE-labeled target cells (**Figure 1E**). In PR8-OVA_II_-infected hosts we found over 20% cytotoxicity (Ag-specific), but little cytotoxicity in PR8-infected hosts. Thus, ThCTL that were cytotoxic to Ag-expressing targets, only developed when effectors re-encountered Ag in the 2^nd^ host.

We also analyzed secretion of canonical Th1 effector cytokines IFNγ and TNFα by lung donor cells. In contrast to ThCTL development that depended on Ag, the donor effectors recovered in the lung did not require Ag recognition to maintain the ability to secrete IFNγ, and TNFα secretion was lost when cognate Ag was present during the effector phase (**Figure 1F-G, Figure S1H**). Thus, we suggest that the components of the program leading to induction of ThCTL phenotypes and functions, but not general Th1 characteristics in the lung, were coordinately driven in 6d effectors by cognate Ag recognition at the effector phase.

### ThCTL development from 6 dpi CD4 effectors does not require CD28 co-stimulation in contrast to T_FH_

CD28 expressed on T cells interacts with CD80/86 on APC during cognate interaction, co-stimulating IL-2 production and initiating proliferation (Watts, 2010). Since we find that CD4 effectors require cognate Ag presentation again to become ThCTL, we analyzed whether the generation of ThCTL from CD4 effectors also requires CD28:CD80/CD86 co-stimulation during this time. As a control, we analyzed generation of T_FH_ (T follicular helper cells) which is another late CD4 effector subset, that is known to require CD28 co-stimulation even during the peak of the effector phase (Linterman et al., 2014). We transferred 6d *in vivo* effectors into WT or CD80/86 deficient PR8-OVA_II_ infection-matched hosts (**Figure 2A**). When hosts lacked the costimulatory ligands for CD28, the recovery of total donor effectors in the lung was significantly decreased (**Figure S2A**), but the proportion of lung donor ThCTL nearly doubled (**Figure 2B**). Donor ThCTL numbers were unchanged (**Figure 2B**), suggesting non-ThCTL were selectively lost. The level of NKG2A/C/E expression on donor ThCTL was also increased in CD80/86 KO compared with WT hosts (**Figure 2B, Figure S2B**), suggesting that the development of full-fledged ThCTL improved without CD28:CD80/86 co-stimulation.

**Figure 2.**
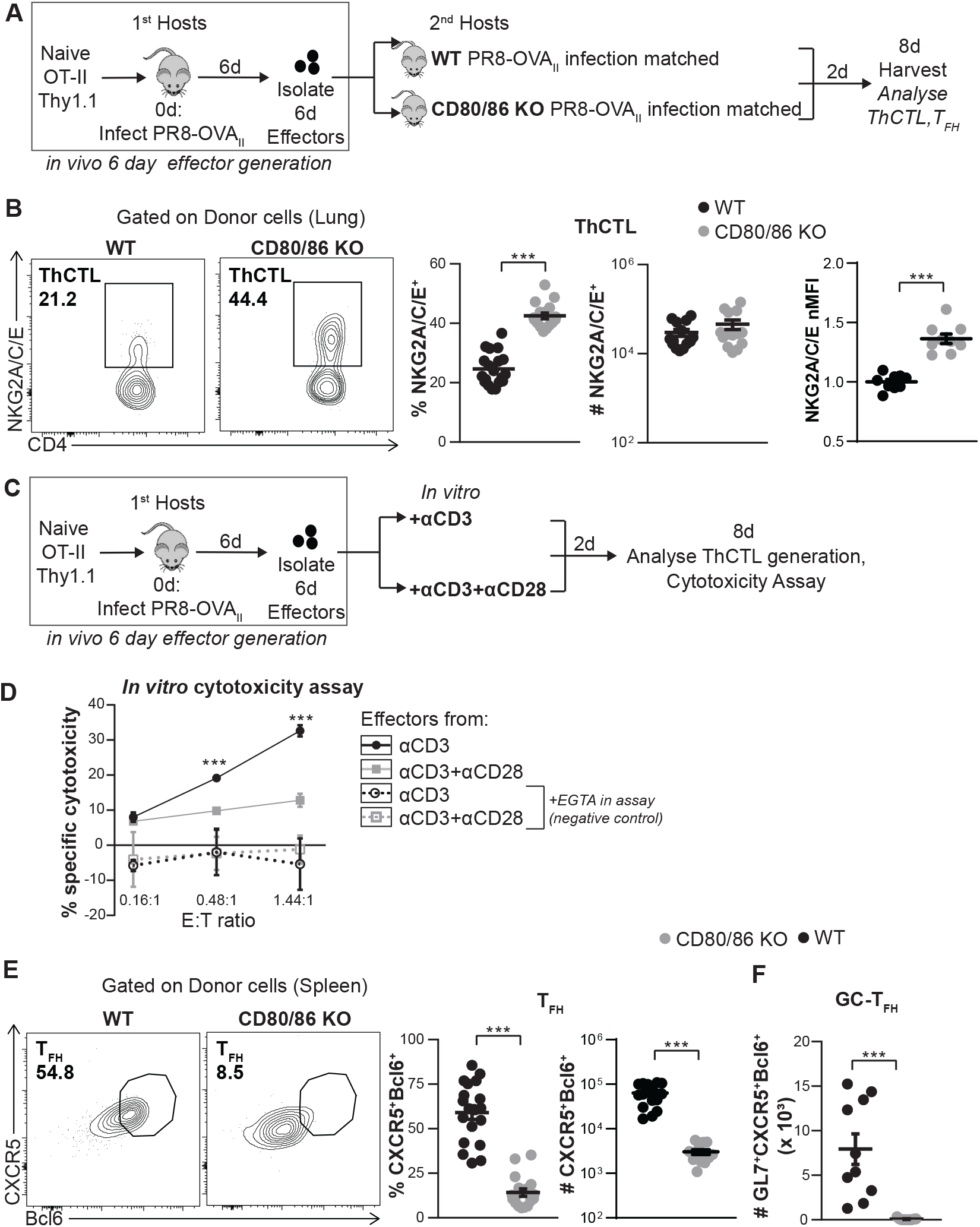
CD4 effectors do not require CD28 co-stimulation to become ThCTL. **(A)** Experimental design for (B, E-F): *In vivo* generated 6d OT-II.Thy1.1^+^ effectors were transferred into 6 dpi PR8-OVA_II_-infected WT or CD80/CD86^-/-^ mice. Spleen, DLN and lungs were harvested at 8 dpi. **(B)** Percentage and number of lung ThCTL (NKG2A/C/E^+^) formed from donor cells. Normalized MFI (nMFI) of NKG2A/C/E expression by ThCTL (n=10-19 per group pooled, 2-4 independent experiments) **(C)** Experimental design: *In vivo* generated 6d OT-II.Thy1.1^+^ effectors were isolated and stimulated with either anti-CD3 alone or anti-CD3 and anti-CD28 *in vitro* to mimic *in vivo* effector phase cognate Ag stimulation. **(D)** Ag specific cytotoxicity of donors generated as in Figure 2C, with anti-CD3 or anti-CD3 + anti-CD28 (Each E:T ratio is assayed in triplicate or single wells for +EGTA conditions, representative of 2 independent experiments). **(E-F)** Experiment done as in Figure 2A. **(E)** Percentage and number of spleen donor T_FH_ (n=14-19 per group pooled, 3-4 independent experiments). **(F)** Number of spleen donor GC-T_FH_ (GL7^+^CXCR5^+^ Bcl6^+^) (n=8-10 per group pooled, 2 independent experiments). Error bars represent s.e.m. Statistical significance determined by two-tailed, unpaired Student’s t-test (* P<0.05, ** P<0.01 and *** P<0.001). See also Fig. S2.

To reproduce this *in vitro*, we cultured *in vivo*-generated 6d effectors for 2d *in vitro* with anti-CD3 vs anti-CD3 plus CD28 (**Figure 2C**). No ThCTL developed without anti-CD3 (**Figure S2C-D**), mimicking the *in vivo* requirement for cognate Ag recognition by CD4 effectors to become ThCTL, as shown in Figure 1. ThCTL developed when effectors were stimulated by CD3 alone and adding CD28 co-stimulation inhibited ThCTL generation (**Figure S2D**). Development of cytotoxic function also depended on anti-CD3 stimulation and was inhibited when CD28 co-stimulation was provided *in vitro* (**Figure 2D**). Together the *in vitro* and *in vivo* results indicate that differentiation of 6d effectors into ThCTL does not require CD28 co-stimulation. The lack of a need for CD28 co-stimulation is also reminiscent of human ThCTL populations that are CD28 negative (Serroukh et al., 2018; van de Berg et al., 2008).

In contrast, in the same experiments, both the proportion and absolute number of donor T_FH_ and GC-T_FH_ in the spleen and DLN were dramatically lower in the CD80/86 KO hosts (**Figure 2E-F, Figure S2E-F**). Thus, while CD4 effectors require CD28 co-stimulation to fully develop into T_FH_ in the spleen and the DLN, they do not require CD28 to sustain or induce further ThCTL generation in the lung. Therefore, while both pathways of specialized CD4 effector development require Ag recognition, the two have distinct co-stimulation requirements.

### Ag presentation by hematopoietic APC effectively induces ThCTL generation from CD4 effectors

A recent study showed that ThCTL generation was significantly reduced when CD11c^+^APC were absent from the initiation of the response (Knudson et al., 2021). Since we show here that 6 dpi effectors need to recognize cognate Ag again to become lung ThCTL, we asked if a specific APC subset is required during this time, to generate ThCTL from already primed 6 dpi effectors. To evaluate the need for DC, we used 3 mouse models: (1) CD11c^Cre^MHC-II^fl/fl^ hosts where CD11c-expressing APC lose MHC-II and thus cannot present Ag (**Figure 3A**) (2) CD11c Tg.H2-Ab1^-/-^ hosts, where the only potential APC subsets expressing MHC-II are CD11c^+^ (**Figure S3A**) and (3) Ag-pulsed DC added as the source of cognate Ag (**Figure S3B**). We found transferring Ag-pulsed DC (OVA_II_/DC) along with 6d effectors into PR8 infection-matched mice was sufficient to drive the CD4 effectors to become ThCTL (**Figure S3B**). There was no defect in donor ThCTL formation when 6 dpi effectors were transferred either into CD11c^Cre^MHC-II^fl/fl^ PR8-OVA_II_ infection-matched mice (**Figure 3C**) or into CD11c Tg.H2-Ab1^-/-^ PR8-OVA_II_ infection-matched mice (**Figure S3A**). Thus, both endogenous and exogenously added MHC-II^+^ DC APC are sufficient to drive ThCTL development, but ThCTL develop from effectors even when CD11c^+^ DC APC are absent, indicating DC are not the only APC that effectively support CD4 effector differentiation to ThCTL.

**Figure 3.**
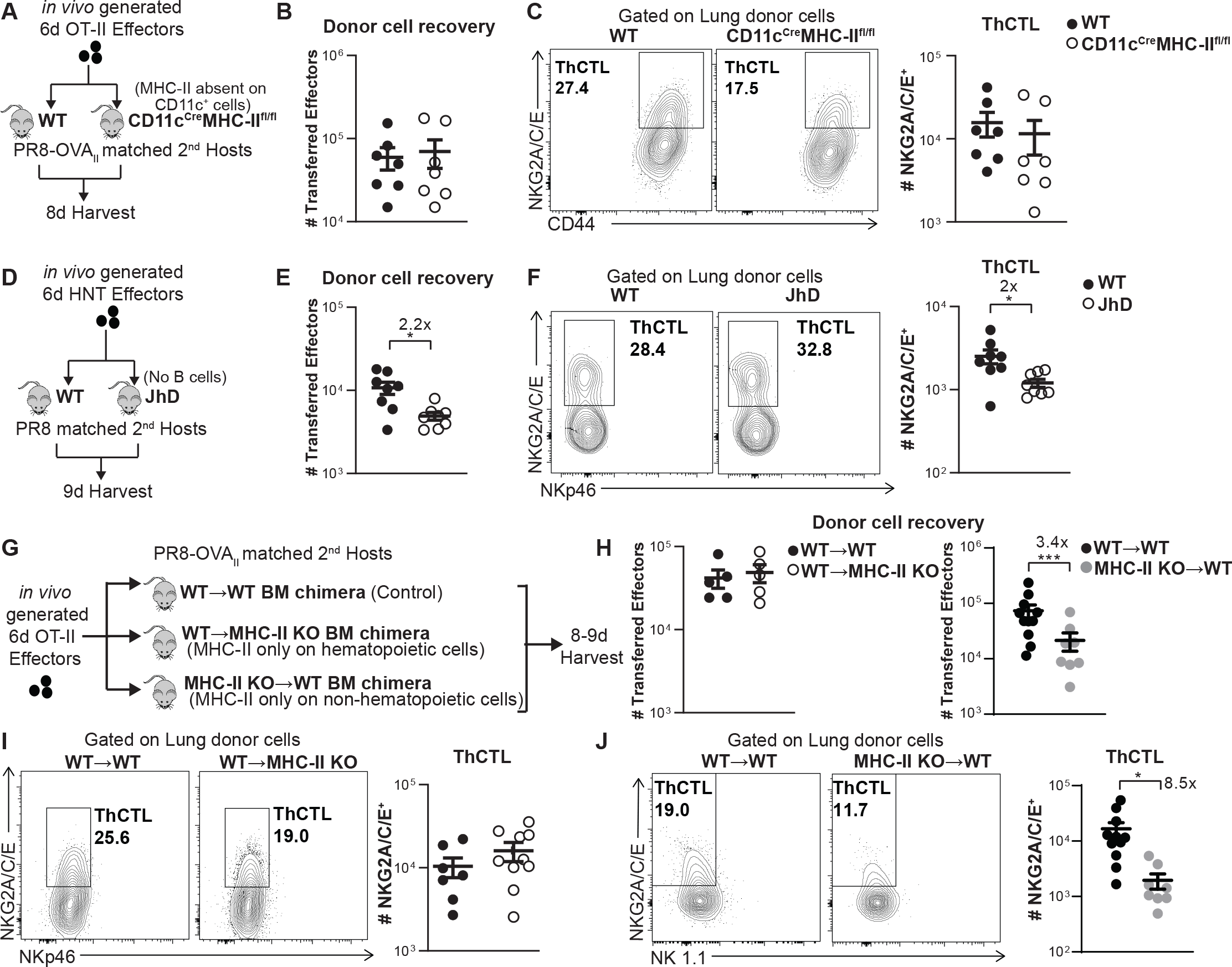
Hematopoietic APC subsets are sufficient to support ThCTL generation from 6d effectors. **(A-C)** *In vivo* generated 6d OT-II.Thy1.1^+^ effectors were transferred into PR8-OVA_II_ infection-matched CD11c^Cre^MHC-II^fl/fl^ hosts where MHC-II Ag presentation on CD11c^+^ APC is absent or into littermate control mice. **(B)** Total number of donor cells in the lung and **(C)** ThCTL generation were analyzed at 8 dpi. (n=7 per group pooled, 2 independent experiments). **(D-F)** *In vivo* generated 6d HNT.Thy1.1^+^ effectors were transferred into PR8 infection-matched JhD hosts that are B cell deficient or into WT control mice. **(E)** Total number of donor cells in the lung and **(F)** ThCTL generation were analyzed at 9 dpi. (n=8 per group pooled, 2 independent experiments). **(G-J)** *In vivo* generated 6d OT-II.Thy1.1^+^ effectors were transferred into PR8-OVA_II_ infection-matched MHC-II KO (H2-Ab1^-/-^) bone marrow chimera mice or into WT chimera control mice. WT→ MHC-II KO chimeras (n=7-8 per group pooled, 3 independent experiments) and MHC-II KO →WT chimeras (n=8-11 per group pooled, 3 independent experiments) were made by transferring WT bone-marrow into MHC-II KO irradiated hosts and vice versa. WT→WT bone marrow chimeras were used as control mice. **(H)** Total number of donor cells in the lung and **(I-J)** ThCTL generation were analyzed at 8-9 dpi. Error bars represent s.e.m. Statistical significance determined by two-tailed, unpaired Student’s t-test (* *P* < 0.05, ** *P* < 0.01 and *** *P* < 0.001). See also Figure S3.

To ask if B cells are required as APC for 6 dpi effectors to become ThCTL in the lung, we used B cell deficient JhD mice. We transferred 6 dpi *in vivo*-generated BALB/c.HNT effectors (specific for influenza A virus hemagglutinin) into PR8 infection-matched BALB/c.JhD vs BALB/c.WT mice (**Figure 3D**). In JhD hosts, there were 2-fold fewer donor effectors in the lung at 9 dpi (**Figure 3E**), but an equivalent proportion of ThCTL generated as in WT hosts, resulting in a 2-fold decrease in number of donor-derived ThCTL (**Figure 3F**). Thus, ThCTL generation from 6 dpi effectors can be induced in the absence of cognate Ag presentation by B cells. To determine if B cells, like DC, were sufficient as APC to drive ThCTL development from effectors, we transferred 6 dpi OT-II effectors with Ag-pulsed B cells (OVA_II_/B) into PR8 infected-matched mice and found ThCTL were generated (**Figure S3C**). Thus, as for DC, B cells are not essential but are sufficient as APC to drive 6 dpi CD4 effector differentiation into ThCTL.

During IAV infection, MHC-II is also upregulated on infected epithelial cells in the lung, a possible source of non-hematopoietic APC (Brown et al., 2012). To determine if non-hematopoietic APC can support ThCTL generation, we transferred 6 dpi effectors into infection-matched bone marrow (BM) chimeras in which MHC II Ag-presentation was restricted to either the hematopoietic compartment [WT→MHC-II KO chimeras] or to the non-hematopoietic compartment [MHC-II KO→WT chimeras] (**Figure 3G**). As expected, there was no defect in ThCTL generation when MHC-II was expressed only by the hematopoietic compartment compared to the controls (**Figure 3I**). On the other hand, when MHC-II was expressed only by non-hematopoietic cells, we found significantly fewer donor ThCTL (8.5-fold decrease) although there were still some generated (**Figure 3J**). This defect was not only due to a decrease in number of total donor effectors in the lung since those were decreased only by 3.4-fold (**Figure 3H**). This indicates that non-hematopoietic APC on their own are inefficient in supporting effector differentiation to ThCTL. We conclude that development of ThCTL from 6 dpi CD4 effectors does not require a unique APC type, that multiple hematopoietic APC including DC and B cells efficiently drive this differentiation and that even non-hematopoietic cells can generate some ThCTL.

### Ag delivery to the lung during the effector phase drives lung ThCTL generation

ThCTL are tissue-resident effectors found in the site of viral replication, which is the lung during influenza virus infection (Marshall et al., 2016). Since 6 dpi effectors isolated from the SLO need to recognize Ag again to become lung ThCTL (Figure 1), we hypothesized that local Ag recognition in the lung may be required.

To evaluate this, we targeted Ag presentation to the lung using intranasal (i.n.) Ag/APC and to the spleen using intrasplenic (i.s.) delivery. We previously showed, using Nur77^GFP^OT-II. Thy1.1 effectors and CD45.1^+^ or GFP^+^ APC, that we can visualize the location of APC and Ag presentation. Transfer of Ag/APC i.n. successfully restricted Ag presentation to the lung, while transfer of Ag/APC i.s restricted Ag presentation to the spleen (Devarajan et al., 2022) (summarized in **Figure S4A**).

We transferred 6 dpi *in vivo*-generated effectors into PR8 infection-matched 2^nd^ hosts with Ag/APC transferred either i.n. or i.s and analyzed ThCTL generation from donor effectors at 9 dpi (**Figure 4A**). Both i.n. and i.s. Ag/APC transfer increased trafficking of total transferred effectors to the lung, compared to the negative control (**Figure 4B**). We compared ThCTL generation after i.n. vs i.s. Ag/APC delivery (**Figure 4C**). Strikingly, only i.n. Ag/APC delivery induced ThCTL. When we used i.s. delivery, few if any ThCTL were generated, suggesting that Ag in the lung is required for lung ThCTL generation.

**Figure 4.**
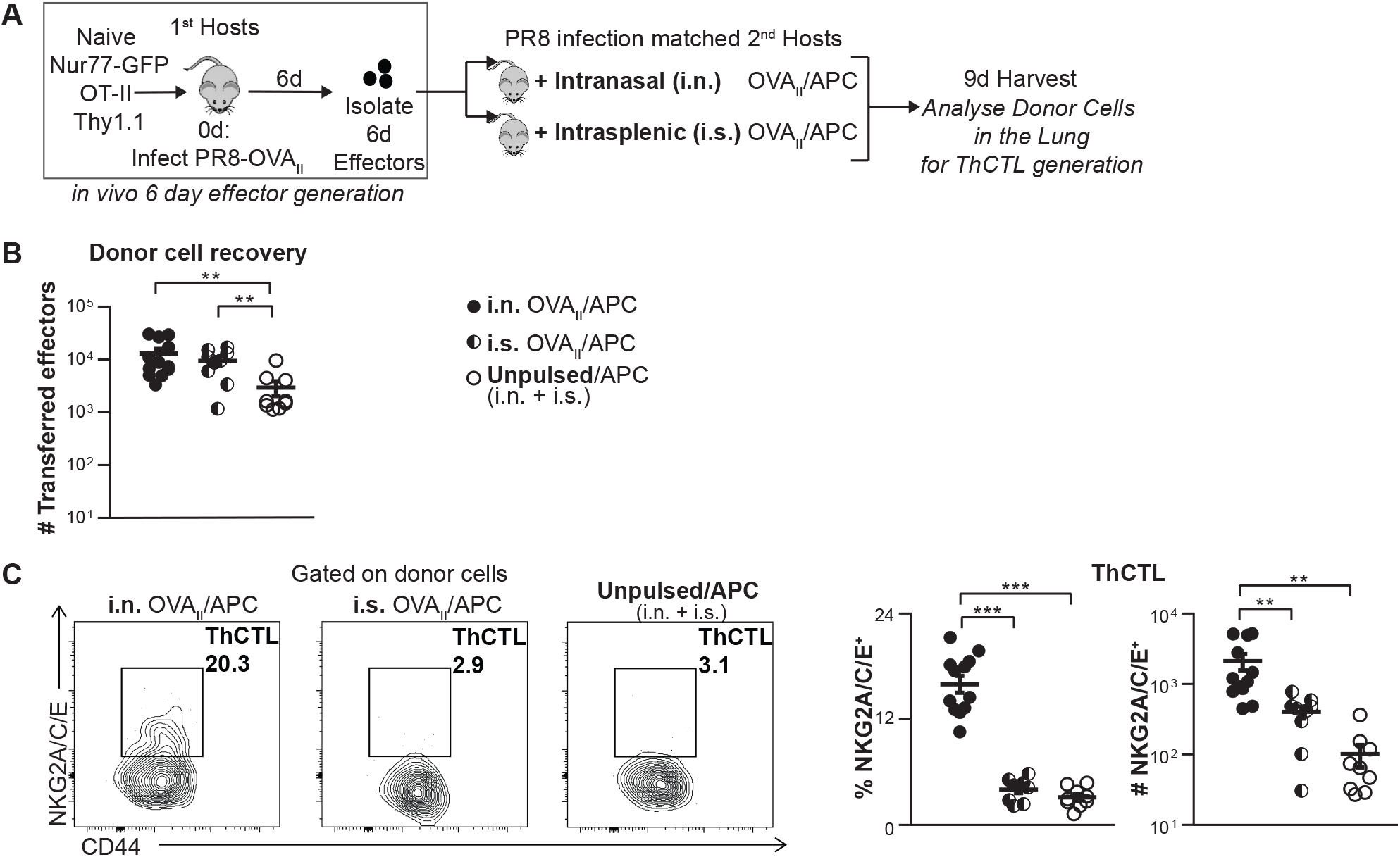
Ag delivery via i.n. and i.s routes, during the effector phase, shows that local Ag presentation in the lung drives ThCTL generation from effectors. **(A)** Experimental design: OVA_II_ peptide pulsed CD45.1^+^ B-cells were used as APC and transferred into PR8 infection-matched hosts 6 dpi either intranasally (i.n.) or intrasplenically (i.s.). Unpulsed APC were transferred both i.n. and i.s. as negative controls. *In vivo* generated 6d OT-II.Nur77^GFP^.Thy1.1^+^effectors were transferred i.v. Mice were harvested 3 dpt and ThCTL generation from donor cells in the lung was analyzed. **(B)** Number of donor effectors recovered with i.n. vs i.s. APC transfer. **(C)** Donor lung ThCTL formation with i.n. vs i.s. APC transfer. (n=9-12 per group pooled, 4 independent experiments) Error bars represent s.e.m. Statistical significance determined by two-tailed, unpaired Student’s t-test (* P < 0.05, ** P < 0.01 and *** P < 0.001). See also Fig S4.

Our previous data show that after intravenous (i.v.) OVA_II_/APC transfer they are found predominantly in the spleen and not in the lung, though donor effectors in the lung express Nur77^GFP^, suggesting that they recognized Ag in the periphery before migrating to the lung (Devarajan et al., 2022) (**Figure S4A**). To test if peripheral Ag presentation was sufficient to drive 6d effectors to ThCTL, we transferred OVA_II_/APC i.v. and compared ThCTL generation to i.n. OVA_II_/APC transfer (**Figure S4B**). OVA_II_/APC transfer i.v., like i.s., increased trafficking of total transferred effectors to the lung, compared to both the negative control and to i.n. transfer (**Figure S4C**). These data (**Figure S4C, Figure 4B**) support the concept that Ag presentation in the spleen enhances pathways that favor migration of CD4 effectors to the lung. In contrast, further development of donor ThCTL in the lung as measured by NKG2A/C/E expression (both percent and MFI) after i.v. transfer, was as low as the negative control (**Figure S4D**). CXCR6 and PD1 expression by the NKG2A/C/E^+^ cells generated with i.v. OVA_II_/APC transfer was also lower compared to those generated with i.n. OVA_II_/APC (**Figure S4E**). This suggests that even if 6d effectors recognize Ag initially in the SLO, and this Ag presentation enhances their migration to the lung, ThCTL develop optimally only when Ag is presented locally in the lung, which is their tissue of residence.

### Infection-induced Type I IFN is required to support the development of ThCTL from 6 dpi CD4 effectors

ThCTL effectors can be pathogenic when dysregulated (Broadley et al., 2017; Juno et al., 2017) as also found recently during COVID infection (Kaneko et al., 2022; Meckiff et al., 2020). We reasoned that ‘danger signals’ from infection might also be required at the effector stage as a mechanism to limit production of lung ThCTL when infection has already been cleared. To analyze whether generation of ThCTL requires infection, independently of Ag presentation, we transferred *in vivo* generated 6d OT-II.Nur77^GFP^.Thy1.1^+^ effectors along with OVA_II_/B cells as APC into IAV infection-matched or uninfected 2^nd^ hosts for 3-4 days (**Figure 5A**). To ensure that the transferred APC present Ag equivalently in both infected and uninfected hosts, we harvested one cohort of each after 14-16hr (**Figure S5A**). We found there was no significant difference either in the number of transferred APC found or in Nur77^GFP^ expression by the donor effectors, in the lung of infected compared to uninfected mice, indicating that we had successfully normalized Ag presentation in both groups of hosts by presenting Ag only on the peptide-pulsed APC (**Figure S5A**).

**Figure 5.**
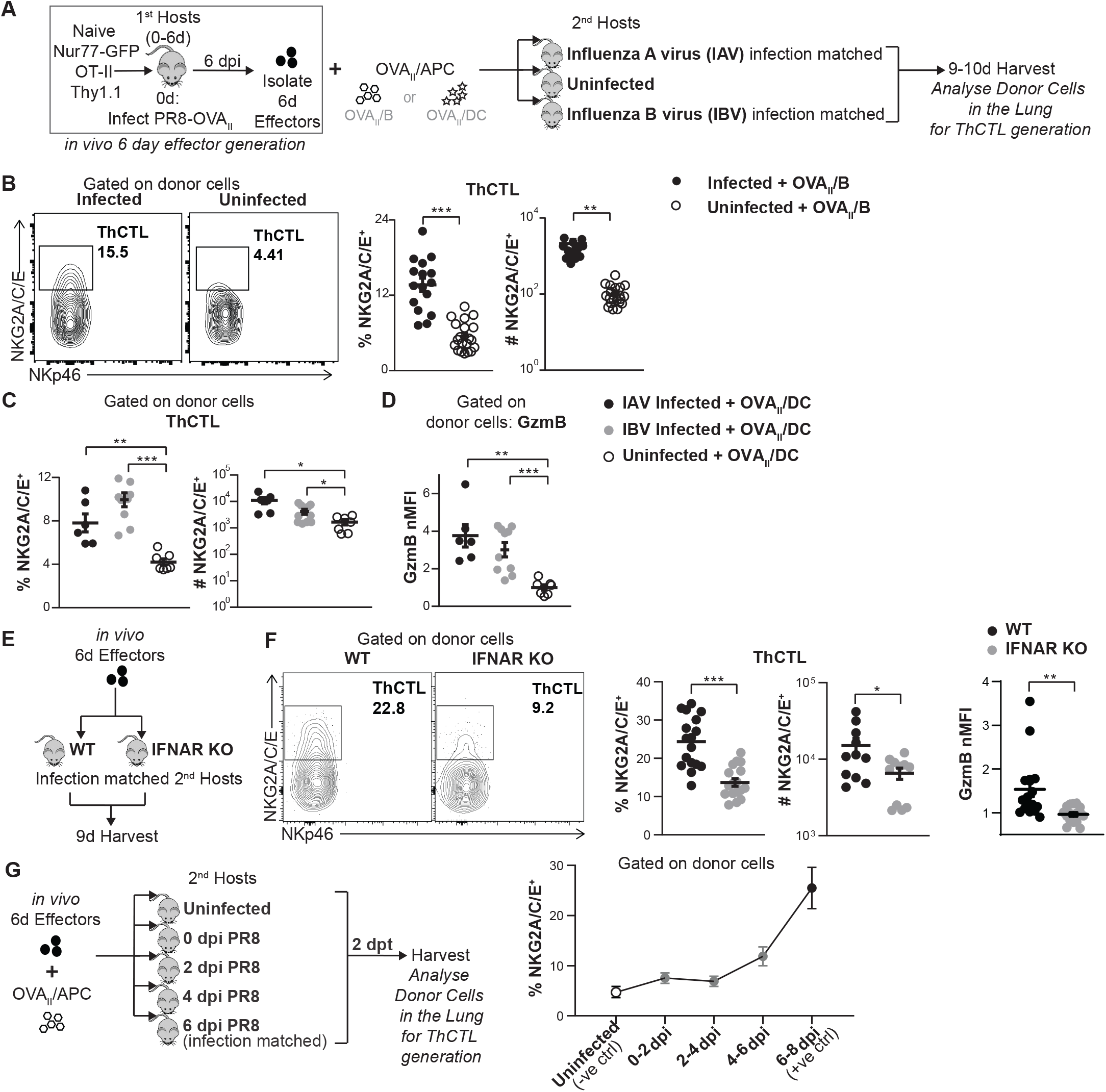
Signals from infection, in addition to Ag recognition, support lung ThCTL generation from effectors. **(A)** *In vivo* generated 6d OT-II.Nur77^GFP^.Thy1.1^+^ effectors were transferred i.v. along with OVA_II_ peptide pulsed B cells (Fig 5B) or DC (Fig 5C-D) that were transferred both i.n. and i.v. into 2^nd^ hosts that were either infection-matched with Influenza A Virus (IAV: PR8), with Influenza B Virus (IBV: B/Ann Arbor/1/66), or that were uninfected. **(B)**6d effectors + OVA_II_/B were transferred into IAV infection-matched vs uninfected hosts and were analyzed by FACS 3-4 dpt for donor lung ThCTL generation (n=16-20 per group pooled from 5 independent experiments). **(C-D)**6d effectors + OVA_II_/DC were transferred into IAV or IBV infection-matched, or uninfected hosts. 4 dpt lungs were analyzed by FACS for **(C)** donor ThCTL generation, **(D)** GzmB normalized MFI of total lung donor cells. (C-D) n=6-10 per group pooled from 2 independent experiments. **(E)***In vivo* generated 6d OT-II.Thy1.1^+^ effectors were transferred i.v. into 2^nd^ wild-type IFNAR KO hosts infection-matched with PR8-OVA_II_ virus. Donor cells in the lung were analyzed 3 dpt **(F)** Lung ThCTL generation from donor cells and donor cell GzmB expression in the lung (n=11-17 per group pooled from 3-4 independent experiments). **(G)***In vivo* generated 6d OT-II.Thy1.1^+^ effectors were transferred i.v. into groups of 2^nd^ hosts that were infected with PR8 at different times before transfer. Equal numbers of OVA_II_ peptide pulsed APC were transferred into all groups of 2^nd^ hosts both i.n. and i.v. T-depleted splenocytes were activated and used as APC. Lung ThCTL generation from donor cells was analyzed 2 dpt. (n=5-7 per group pooled from 2 independent experiments). Error bars represent s.e.m. Statistical significance determined by two-tailed, unpaired Student’s t-test (* *P*< 0.05, ** *P*< 0.01 and *** *P*< 0.001). See also Fig. S5.

We assessed donor ThCTL generation at 3-4 dpt. Lung ThCTL recovery in infection-matched positive controls was markedly higher than in uninfected mice (**Figure 5B**). Donor cells in the infected host lung expressed increased GzmB, increased PD1 and decreased CXCR3 compared to uninfected hosts, each of which are associated with ThCTL (**Figure S5C**). To investigate whether the impact of infection was linked to Ag presentation, we instead infected with Influenza B virus (IBV) (**Figure S5D-G**). IBV shares no CD4 T cell epitopes with IAV, therefore IBV-infected 2^nd^ hosts only provide infection-generated signals but do not present cognate Ag. We supplied cognate Ag presentation with OVA_II_/B cell APC. ThCTL generation was highly increased in IBV infection-matched compared to uninfected mice (**Figure S5E**). GzmB expression by lung donor cells was also drastically increased by IBV infection compared to uninfected hosts (**Figure S5E**). To evaluate this requirement for infection with another APC subset, we repeated the experiment using DC as the source of OVA_II_/APC (**Figure 5C-D**). Again, infection was needed for optimum generation of ThCTL and either IAV or IBV support this. Infection also enhanced GzmB expression and associated ThCTL phenotypes (**Figure 5D, S5G**). Thus, infection plays a clear role in inducing ThCTL, distinct and in addition to its role in providing local Ag presentation.

Since induction of Type I IFN is classically associated with viral infections, we tested if infection-induced Type I IFN in the host environment, plays a role in ThCTL generation from 6 dpi effectors. We transferred 6 dpi *in vivo*-generated WT effectors into infection-matched IFNAR KO hosts and analyzed their development into lung ThCTL (**Figure 5E**). Fewer donor cells became ThCTL and expressed lower GzmB in IFNAR KO compared to those in WT lungs (**Figure 5F**). However, there was no decrease in total donor effectors recovered from the lung (**Figure S5H**). This suggests a role for Type I IFN signals in driving induction of ThCTL. In this model the host mice lacked IFNAR while the transferred 6 dpi WT effectors expressed IFNAR, suggesting that Type I IFN played an indirect role by inducing a factor in host cells that was required for ThCTL generation.

To evaluate when infection-induced factors may play a role, we asked when after infection the signals supporting ThCTL development were optimal. We transferred 6 dpi effectors with OVA_II_/APC into 0, 2, 4 and 6 dpi PR8-infected hosts and harvested lungs from each group after 2 dpt (**Figure 5G**). Lung ThCTL were most efficiently generated in mice infected for 6-8 days (**Figure 5G**, **Figure S5I**) suggesting that type I IFN induces a factor that is required for ThCTL generation, which is at sufficient levels only after 6 dpi.

### IL-15 synergizes with local cognate-Ag recognition in the lung to drive ThCTL generation from 6 dpi effectors

Since the factor(s) supplied by infection that are required for ThCTL generation in addition to Ag presentation is best induced after 6 dpi, we analyzed pathways that were upregulated in lung CD4 effectors at that time. We performed scRNAseq analysis of influenza-infected lungs at 6 dpi and compared naïve CD4 clusters with activated CD4 clusters. The differentially expressed genes showed an enrichment for a Hobit-regulated pathway that is induced by IL-15 (Mackay et al., 2016; Mackay et al., 2015) (**Figure 6A**). As Type I IFN induced a factor(s) that could drive the generation of lung ThCTL (Figure 5E-F), we asked if Type I IFN might act by inducing IL-15. We quantified levels of IL-15 in IFNAR KO vs WT lungs at various dpi. In mice without IFNAR, there was a reduction in IL-15 levels in the bronchoalveolar lavage (BAL) after 5 dpi, as determined by qPCR (**Figure 6B**). IL-15 is predominantly transpresented by the IL-15Rα on the surface of cells (Nolz and Richer, 2020), so only minimal quantities of IL-15/IL-15R complex are secreted that can be quantified by ELISA. There was also a reduction in the secreted IL-15/IL-15R complex in the BAL fluid (BALF) of IFNAR KO mice (**Figure 6B**). This is in line with previous studies that show that Type I IFN induces IL-15 during infection (Colpitts et al., 2012; Colpitts et al., 2013; Mattei et al., 2001). scRNAseq of whole 6 dpi lungs showed that IL-15 and IL-15Rα were expressed mainly by subsets of macrophages, neutrophils, endothelial cells and fibroblasts, suggesting that these subsets may have the potential to transpresent IL-15/IL15-Rα (**Figure S6A**).

**Figure 6.**
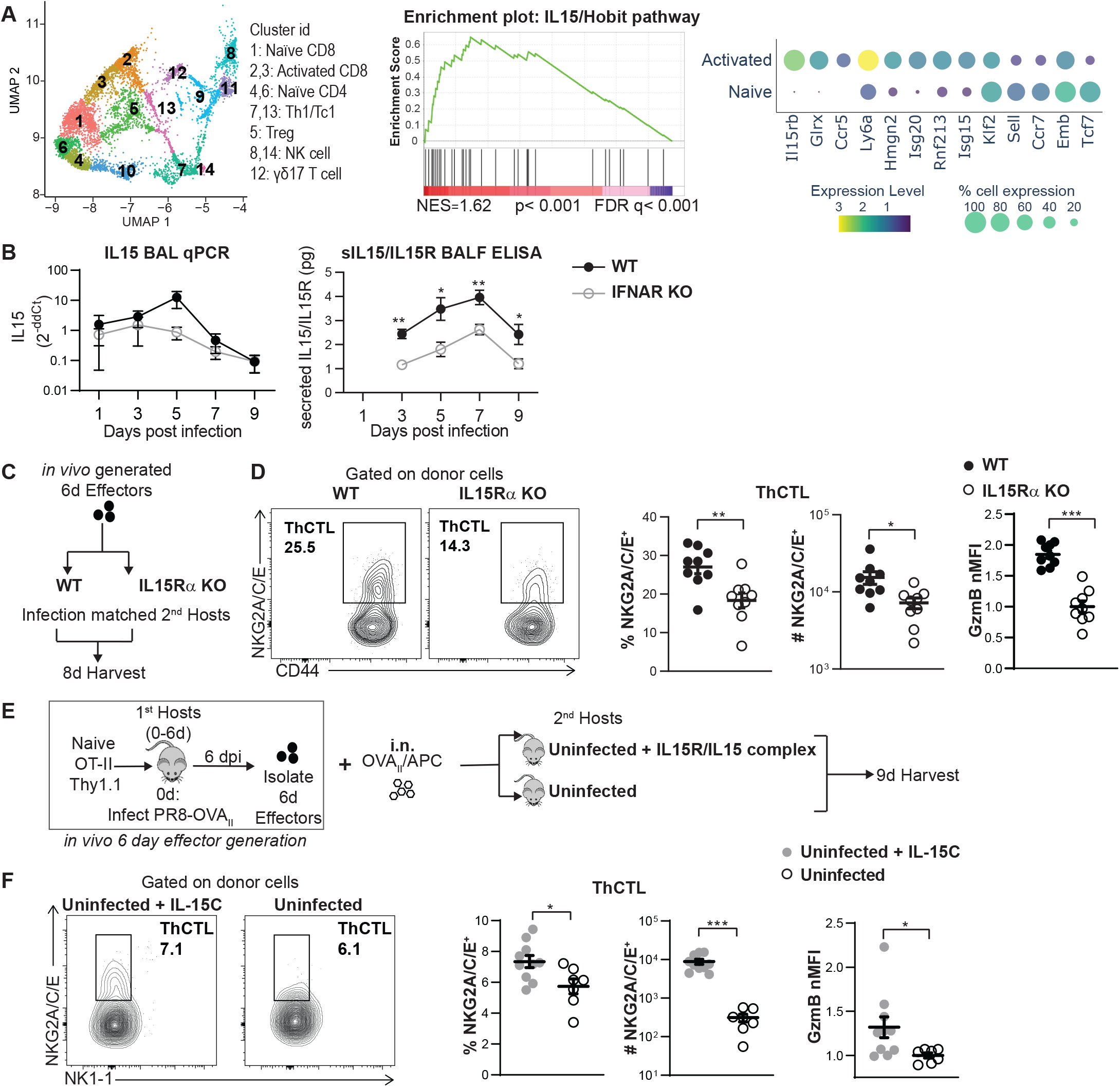
CD4 effectors require IL-15 to become lung resident ThCTL effectors. **(A)** Influenza infected lungs were analyzed by scRNAseq at 6 dpi. Naïve CD4^+^ cells in clusters 4&6 were compared with activated CD4^+^ cells in clusters 7&13. GSEA was used to test if a IL-15 induced Hobit pathway (Mackay et al, 2016) was enriched in lung CD4 effectors. Expression of a panel of IL-15/Hobit regulated genes are plotted. Size of dots represent the percent of cells in the population expressing the gene. Color of the dots represents the normalized expression value of the gene. **(B)** IFNAR KO and WT control mice were infected with PR8 virus. Their bronchoalveolar lavage (BAL) was collected at various dpi. IL15 gene expression in the BAL cells was analyzed by qPCR. Secreted IL-15/IL-15R levels in the BALF was analyzed by ELISA. (n=5-11 per group pooled, 2-4 independent experiments). **(C)** *In vivo* generated 6d OT-II.Thy1.1^+^ effectors were transferred into PR8-OVA_II_ infection-matched IL-15Rα KO hosts or into control mice. Donor cells in the lung were analyzed 2 dpt **(D)** Donor ThCTL (NKG2A/C/E^+^) generation and donor GzmB nMFI expression was analyzed in the lung (n=9-10 per group pooled, 2 independent experiments). **(E)** *In vivo* generated 6d OT-II.Thy1.1^+^ effectors were transferred into two groups of uninfected hosts along with OVA_II_/APC that were transferred i.n. T-depleted splenocytes were activated and used as APC. One group was treated i.n. with IL-15/IL-15Rα complex starting the day before transfer to the day before harvest. Control group was treated i.n. with PBS. **(F)** ThCTL generation from donor cells and donor GzmB expression was analyzed 3 dpt in the lung. (n=8-10 per group pooled, 2 independent experiments). Error bars represent s.e.m. Statistical significance determined by two-tailed, unpaired Student’s t-test (* *P* < 0.05, ** *P* < 0.01 and *** *P* < 0.001). See also Fig. S6.

To specifically assess the role of IL-15 in driving ThCTL generation, we transferred 6 dpi effectors into infection-matched IL-15Rα KO hosts (**Figure 6C**), that cannot transpresent IL-15 to CD4 effectors. Indeed, fewer donor ThCTL were generated from 6 dpi effectors in the IL-15Rα KO hosts (**Figure 6D**), suggesting a role for IL-15 in ThCTL generation. To confirm that IL-15 synergizes with local Ag presentation in the lung to generate ThCTL from 6 dpi peripheral effectors, we transferred 6 dpi *in vivo-generated* effectors along with i.n. OVA_II_/APC into uninfected hosts, with or without IL-15/IL-15Rα complex administered i.n. (**Figure 6E**). Many more ThCTL were generated from 6 dpi effectors in uninfected hosts when intranasal IL-15 complexes were provided (**Figure 6F**), and they were generated to levels comparable to those seen during influenza infection (**Figure S6E**). Thus, infection-induced Type I IFN induces IL-15 production in the lung which in turn supports lung ThCTL generation from 6 dpi SLO CD4 effectors.

## DISCUSSION

ThCTL contribute to viral clearance using perforin-dependent, MHC-II restricted cytotoxicity and synergize with B cell responses to enhance recovery from infections (Brown et al., 2006; Brown et al., 2012; Fang et al., 2012; Marshall et al., 2016). We have shown that lung-resident ThCTL help clear influenza virus and recover from primary influenza infections (Brown et al., 2006; Brown et al., 2012; Marshall et al., 2016). Lung-resident ThCTL have also recently been documented in lungs of COVID patients (Kaneko et al., 2022). There are several discrete subsets of cytotoxic CD4 T cells that are defined by their location including circulating cytotoxic CD4, CD4 IEL and tissue-resident ThCTL (Juno et al., 2017; Marshall et al., 2016). The requirements for the generation of ThCTL, that are bona fide CD4 T cells expressing ThPOK, and that are induced during viral infections is largely undefined in vivo. We show that influenza infection-induced ThCTL are generated from peripheral 6 dpi effectors that traffic to the lung where they must once again recognize local cognate Ag on MHC-II^+^APC. In addition, full-fledged ThCTL development requires the infection to induce type I IFN, that acts on host cells to produce IL-15, which drives ThCTL generation from CD4 effectors. Our results also suggest that tissue residency in CD4 effectors is established by one or more of these signals. Thus, ThCTL development is tightly regulated and depends on continuing infection at the effector stage.

We previously established that cognate Ag recognition by CD4 effectors and autocrine IL-2 induction is required for generation of protective CD4 memory, and that these signals are required by CD4 effectors at 6-8 dpi (Bautista et al., 2016; McKinstry et al., 2014; Swain et al., 2021). Recently we showed that T_FH_ development during IAV infection required cognate Ag recognition by CD4 effectors in the SLO (Devarajan et al., 2022), and signals from CD28 co-stimulation. Here we identify the key signals for CD4 effectors to transition into ThCTL. This suggests that there is an effector checkpoint, at which CD4 effectors must receive multiple signals from infection unique to each fate, to progress into each of these specialized CD4 subsets. This effector checkpoint coincides *in situ* with the peak of the CD4 effector response, which is followed by rapid contraction, supporting the concept that CD4 effectors express a default program of apoptosis which they avoid only when they recognize Ag (Bautista et al., 2016; McKinstry et al., 2014). These studies together with our findings here indicate that distinct sets of signals, during the effector checkpoint, concurrently drive alternate fates for CD4 effectors including T_FH_, ThCTL and CD4 memory. Thus, even if CD4 effectors are pre-committed to certain fates early on, they need further signals at this effector checkpoint to realize their potential and complete their development into these subsets. Our results here imply that the majority of vaccines, that are known to present antigen and pathogen recognition only for a few days, will lead to little generation of ThCTL or memory derived from ThCTL. Since many viruses evade CD8-mediated cytotoxicity by downregulating MHC-I, lack of CD4 driven cytotoxicity by ThCTL that recognize MHC-II presented antigens, is likely to lead to poor vaccine-induced protection.

One pathway-specific signal is CD28 co-stimulation that is required for T_FH_, but not for ThCTL generation (Figure 2) which it may limit. This finding also fits with studies that indicate human ThCTL are CD28^-^ (van de Berg et al., 2008). The opposing CD28 requirement for T_FH_ vs ThCTL generation mirrors their opposing requirements via the Bcl6-Blimp axis (Donnarumma et al., 2016; Johnston et al., 2009; Marshall et al., 2016), suggesting these two specialized CD4 effector subsets follow opposing axes of development, though they develop concurrently during infection.

IL-15 is well-known as a homeostatic and survival factor, but has also been proposed to be a danger signal that communicates an altered tissue state to the immune system, thus licensing more potent effector functions (Jabri and Abadie, 2015). This seems to be the case during ThCTL development, where IL-15 peaking at 5-7 dpi in the infected lung tissue, is required for CD4 effectors to pass the effector checkpoint and become cytotoxic (Figure 6). In NK cells and CD8 T cells, IL-15 induces the expression of cytolytic pore forming perforin and death-inducing granzymes (Jabri and Abadie, 2015; Nolz and Richer, 2020), compatible with our finding here that IL-15 enhances ThCTL in the lung during influenza infection. Mackay et al have shown that IL-15 also induces CD8 T_RM_ via a Hobit/Blimp dependent pathway (Mackay et al., 2016; Mackay et al., 2015). Our scRNAseq data indicates that a Hobit pathway is indeed expressed in lung CD4 effectors along with downregulation of genes such as Klf2, Ccr7, Sell and Tcf7 that result in tissue retention (Mackay et al., 2016). This suggests IL-15 may act by a similar mechanistic pathway through Hobit to induce lung-residence in ThCTL effectors, a hypothesis that needs to be further explored. Thus IL-15 may be playing a dual role inducing both cytotoxicity as well as tissue residence in lung ThCTL during influenza infection.

While various MHC-II^+^ APC, including DC and B cells, could support donor effectors to develop into ThCTL (Figure 3), ThCTL developed only when Ag/APC were delivered to the site of residence in the lung (Figure 4). Recently, Ag presentation in tissues has been implicated in development of both T and B resident memory subsets (Allie et al., 2019; Khan et al., 2016; McMaster et al., 2018; Takamura et al., 2016). Our data suggest that local Ag presentation in the lung along with IL-15 likely establishes tissue residency in ThCTL effectors during the effector checkpoint, and we suggest this most likely is carried over when they become resting memory.

Our results show that the generation of lung ThCTL requires infection to continue into the effector phase (Figure 5–6) and that the infection-generated signals that are required in addition to local Ag presentation, are only present optimally after 6 dpi. A recent study using a mousepox model of Ectromelia infection also showed that ThCTL do not develop if virus is cleared early (Knudson et al., 2021). We find few ThCTL develop when infection, as sensed by Type I IFN induced IL-15, is absent at this effector checkpoint. Type I IFNs are induced by many pathogens and are considered a proxy for ongoing infection. Type I IFN induces IL-15 in myeloid cells in response to viral infection (Colpitts et al., 2012; Colpitts et al., 2013; Mattei et al., 2001). We postulate that requiring simultaneous local Ag recognition and infection-generated type I IFN and IL-15, acts as a safeguard to limit ThCTL generation only to those situations where there is still an ongoing threat at the effector stage. This would limit unnecessary, potentially harmful responses. Indeed, several reports describe exaggerated ThCTL responses in certain autoimmune diseases and infections (Broadley et al., 2017; Juno et al., 2017) where continuous Ag and inflammation persist, including severe COVID infection (Kaneko et al., 2022; Meckiff et al., 2020). Our findings here also suggest that interventions such as blocking IL-15 or local Ag presentation, may attenuate exaggerated ThCTL responses seen during COVID infections.

ThCTL participate in viral clearance (Marshall and Swain, 2011), so an optimal and well-regulated ThCTL response is likely needed for a successful response to influenza and other respiratory viruses. In our studies ThCTL protected against lethal influenza infection by synergizing with humoral responses via a perforin-dependent mechanism (Brown et al., 2006). Critically, their cytotoxicity is not sensitive to viral evasion mechanisms that downregulate MHC-I and thus impede CD8 dependent cytotoxicity (Marshall and Swain, 2011; Soghoian and Streeck, 2010). With cytotoxic CD4 being linked to sustained responses with CAR-T therapy (Melenhorst et al., 2022), it is likely that ThCTL play important roles in anti-tumor immunity, particularly in tumor tissues that downregulate MHC-I to evade immune responses. Thus, strategies to better induce ThCTL effectors in tumor sites could likely benefit by a design that provides prolonged Ag and IL-15 signals in tumor sites.

Coupled with our earlier studies of CD4 memory (Bautista et al., 2016; McKinstry et al., 2014) and T_FH_ generation (Devarajan et al., 2022), our results here support a new paradigm in which a set of critical fate decisions occur at the CD4 effector checkpoint to coordinately support generation of multiple alternate fates: CD4 memory, T_FH_ and ThCTL. We suggest that most effective immunity will develop only when acute infection or novel vaccines provide the effector checkpoint signals identified here, at the right time and in the relevant local tissue sites, so as to drive robust tissue-resident effectors and subsequent CD4 memory resulting in stronger protection against future infections (Devarajan et al., 2016).

## ACKOWLEDGEMENTS

We would like to thank Dr. Laura Mackay and Dr. Luke Gandolfo (University of Melbourne) for their assistance acquiring the IL15/Hobit regulated dataset; Dr. Niranj Chandrasekaran (Broad Institute) for help analyzing the scRNAseq data; and Swain Lab members – Dr. Richard Dutton, Dr. Kai McKinstry, Dr. Michael Jones, Dr. Esteban Rozen, Michael Perkins and Jialing Liang for their intellectual input and assistance with experiments.

## AUTHOR CONTRIBUTIONS

P.D. and S.L.S. wrote the manuscript with assistance from other authors. P.D., A.M.V. and S.L.S. conceived the project and designed experiments. P.D., A.M.V. and N.J.S. performed and analyzed experiments with assistance from C.H.C., O.K.U. and B.L.B. A.M.V. primarily performed and analyzed experiments for Figures 1–2 and associated supplementary figures. P.D. primarily performed and analyzed experiments for Figure 3–6 and associated supplementary figures. N.J.S performed the scRNAseq. K.A.K. performed intrasplenic transfers with assistance from P.D. All authors have read and approved the submitted version.

## MATERIALS AND METHODS

### Mice

C57Bl/6 (B6), B6.CD45.1, B6.Thy1.1, B6.Nr4a1^eGFP^ (Nur77^GFP^), B6.CD80/CD86 KO, IL-15Ra KO, CD11c-Cre-GFP, I-AB-flox and B6.MHC II^-^ were obtained from the Jackson Laboratory. B6.TCRα/β KO mice were obtained from R.Welsh (UMass Chan), Y-linked B6.OT-II mice from L.Bradley (The Scripps Research Institute, La Jolla, CA) and were originally published by F.Carbone’s group (Barnden et al., 1998), BALB/c.HNT from D. Lo (Scott et al., 1994) (The Scripps Research Institute, La Jolla, CA), IFNAR KO from J.Sprent (Garvan Institute), CD11cTg.H2-Ab1^-/-^ mice from T.Laufer originally and all have been bred and maintained at the UMass Chan animal facility. Mice were at least 8 weeks old prior to use. Experimental animal procedures were done in accordance with UMass CHan Animal Care and Use Committee guidelines.

### Virus stocks and infections

Influenza A viruses (IAV) A/Puerto Rico/8/34 (PR8), originally from St. Jude Children’s Hospital, and A/PR8-OVA_II_, kindly provided by Dr. Peter Doherty, were grown and maintained at the Trudeau Institute. Influenza B virus (IBV) B/Ann Arbor/1/86, kindly provided by Dr. Suzanne Epstein (FDA, CBER), was grown and maintained at the Trudeau Institute. Mice were anesthetized with isoflurane (Piramal Healthcare) or with Ketamine/Xylazine (at a dose of 100/10mg/kg by i.p. injection) and were infected intranasally with influenza virus corresponding to a 0.2-0.3 LD_50_ dose in 50 uL of PBS.

**Bone marrow chimera mice** were generated as described previously (Devarajan et al., 2022)

### *In vivo* day 6 effector generation and transfer/*in vitro* culture

*In vivo* generated 6d CD4 T cell effectors were routinely obtained as described previously (Bautista et al., 2016). Briefly, cells from lymph nodes and spleens of naïve OT-II or HNT transgenic mice were enriched for naïve cells by percoll gradients and CD4 T cells isolated by CD4 positive selection (Miltenyi Biotec) or using a CD4 naïve positive selection kit (Miltenyi Biotec). Naïve CD4 T cells were adoptively transferred into mice (1^st^ hosts), which were then infected with IAV (PR8 or PR8-OVA_II_). On day 6 post infection, the lung draining lymph nodes (DLN) and spleens were harvested and donor T cells were isolated using MACS (Miltenyi Biotec) based on their congenic marker (CD90.1). Immediately after isolation, 1-2×10^6^ *in vivo* generated 6d CD4 effectors were adoptively transferred intravenously (i.v.) into host mice (2^nd^ hosts).

*In vivo* generated 6d CD4 effectors were also cultured *in vitro* for 2 days by stimulating with plate bound anti-CD3 (2C11, 0.5ug/ml) or anti-CD3 and anti-CD28 (37.51, 20ug/mL) in T cell media (RPMI 1640 supplemented with 7.5% fetal bovine serum, 0.36mM L-glutamine, 50 uM 2-mercaptoethanol, 162 IU penicillin, 162 ug/ml streptomycin and 10mM HEPES).

### *In vitro* APC culture and activation

BMDC (bone marrow derived dendritic cells) (Bautista et al., 2016; Brahmakshatriya et al., 2017) and activated B cell (Bautista et al., 2016) generation was done as described previously. When a specific APC subset was not required, APC were generated using naïve splenocytes T-depleted using CD90.2 negative selection (Miltenyi Biotec) and cultured *in vitro* for 2 days with 10ng/mL LPS and 10ng/mL dextran sulfate.

### *In vivo* APC delivery

To deliver Ag/APC (BMDC or activated B cells), APC were pulsed with 10μM OVA_323–339_ (OVA_II_) peptide (New England Peptide) or no peptide as a negative control (unpulsed APC) for 1 hour at 37°C with shaking. APC were washed and administered either intravenously (i.v.) in 200uL PBS, intranasally (i.n.) in 50uL PBS, or intrasplenically (i.s.) in 10uL PBS. 0.25-1×10^6^ BMDC or 1×10^6^ B cells were transferred i.v., 0.5-2×10^6^ BMDC or 1-2×10^6^ B cells were transferred i.n. and 0.5-1×10^6^ B cells were transferred i.s.

Intrasplenic transfer of APC was performed as described previously (Devarajan et al., 2022).

### T cell functional assays

*In vivo* and *in vitro* cytotoxicity was performed as previously described (Marshall et al., 2016). Briefly, for *in vivo* cytotoxicity, T depleted splenocytes were stained with either 1uM or 0.4uM of CFSE denoting target (0.4uM) or bystander (1uM) cells. Target cells were pulsed with OVA_II_ peptide for 1 hour at 37°C. Both populations were washed twice in PBS and adoptively transferred into host mice. 18 hours later, the spleens of host mice were harvested and the number of target and bystander cells were quantified by flow cytometry. Specific killing was calculated as: 100 x (1- (live targets/live bystanders)) normalized to the ratio found in control mice. For *in vitro* cytotoxicity, targets were activated B cells that were labeled as above using CellTrace Violet (Invitrogen). Effectors and targets were co-cultured in 96 U bottom plates in T cell medium at 37°C 5% CO_2_ for 4 hours. Plates were washed and stained for cell viability using Annexin V and 7-AAD (Invitrogen) or live/dead amine dyes (Invitrogen). Ag specific cytotoxicity was calculated as: 100 x (1- (live targets/live bystanders)) normalized to the ratio found in control wells with no effector cells. T cell degranulation and cytokine production was measured by *ex vivo* stimulation with plate bound 0.5ug/mL anti-CD3 and 20ug/mL anti-CD28 or with 10ng/mL PMA and 500ng/mL Ionomycin for 4 hours at 37°C, 5% CO_2_ with brefeldin A (10ug/ml). T cell degranulation was also measured simultaneously with the addition of anti-CD107a PE (Biolegend, 1:200), and monensin (BD GolgiStop, according to manufacturer’s protocol) at the beginning of the culture. Cells were harvested and stained for intracellular cytokines.

### Flow cytometry

Cells were processed, stained and analyzed flow cytometry as previously described in detail (Devarajan et al., 2022). Surface antibodies used: anti-CD4 (GK1.5), CD19 (6D5), CD44 (IM7), CD90.1 (OX-7 and HIS51), CD107a (1D4B), CD183 (CXCR3, CXCR3-173), CD185 (CXCR5, SPRCL5), CD186 (CXCR6, SA051D1), CD335 (NKp46, 29A1.4), GL-7, NK1.1 (PK136), and NKG2A/C/E (20d5). Binding to P-selectin was measured by incubating with P-selectin IgG Fusion protein (BD Bioscience), washed and detected with fluorochrome conjugated secondary goat anti-human antibodies (Jackson ImmunoResearch). Subsequent staining for cytokines using the following antibodies: anti-IFNγ (XMG1.2), anti-TNFα (MP6-XT22). GzmB was stained intracellulary directly *ex vivo* using anti-GzmB (GB11). Intracellular Bcl-6 stained with anti-Bcl-6 (K112-91). Antibodies were obtained from eBioscience, Biolegend, or BD Bioscience.

### IL-15 complex treatment

Recombinant Mouse IL-15R alpha Fc Chimera Protein (R&D Systems) and recombinant Mouse IL-15 (Tonbo Biosciences) were suspended and mixed in PBS, then incubated in a 37^0^C water bath for 20-30 minutes (Stoklasek et al., 2006). The mix was immediately placed in ice and mice were promptly treated. Each mouse was anesthetized (Ketamine/Xylazine 100/10mg/kg i.p.) and received 0.75ug of IL-15 precomplexed with 4μg of IL-15Rα-Fc in 50 μl of PBS i.n.

### IL15 qPCR

Cells from the bronchoalveolar lavage (BAL) of mice were resuspend in Trizol and RNA extracted as per manufacturer protocol. RNA was DNAse treated, and reverse transcribed into cDNA using SuperScript II (Invitrogen). Duplex Quantitative PCR was performed using the TaqMan Universal PCR Mastermix (Applied Biosystems) to amplify IL15 gene and 18SrRNA housekeeping gene, with the Bio-Rad CFX96 Real-Time PCR Detection System. Pre-Developed TaqMan primers (Applied Biosystems) for IL15-FAM (Mm00434210_m1) were used multiplexed with 18SrRNA-VIC/MGB housekeeping control primers using the recommended cycling conditions. IL15 gene expression for samples was normalized to that in mock infected lungs.

**IL-15/IL-15R Complex ELISA** was performed using a kit (Invitrogen Cat# 88-7125) following manufacturer recommended protocol. ELISA plates were incubated with BAL fluid (BALF) samples at 4°C overnight. Plates were read with a Spectramax190 microplate reader.

### Cell isolation, library preparation, and data analysis for single cell RNA-Seq

10-week-old male C57BL/6J mice bred for one generation in the UMMS animal facility, from animals purchased from Jackson Laboratory were infected with PR8 as described above. At 6 days post infection, 12 infected and 4 uninfected, animals were sacrificed and perfused with 10ml cold PBS. Lungs were collected and single cell suspensions were prepared using the Miltenyi Biotec Lung Dissociation Kit (130-095-927) with the gentleMACS Octo Dissociator (130-095-937) following the manufacturer instructions. Cells were loaded onto the Chromium Controller (10x Genomics) and single-cell RNA-seq libraries were prepared using the Chromium Single Cell 3□ Reagent Kits v3 (10x Genomics) following the standard manufacturer protocols (one uninfected animal sample was excluded due to reagent clog during cell partitioning resulting). The libraries were sequenced on an Illumina NovaSeq S4, and resulting sequencing data were processed using Cell Ranger (10x Genomics, v3.1.0) within the DolphinNext Cell Ranger Pipeline (Revision 2) (Yukselen et al., 2020). Reads were aligned to both the mm10 and PR8 genomes and resulting count matrices were loaded into Monocle 3 (v1.0.0) for analysis (Cao et al., 2019; Trapnell et al., 2014). Cells expressing fewer than 100 genes or with high mitochondrial content were discarded. Normalization and PCA were performed using the preprocess_cds function, UMAP was performed to visualize the cells using the reduce_dimensions function on the top 100 principal components, and clusters were identified using the cluster_cells function using default settings. Marker genes for each cluster identified using the top_markers function were used to identify cell types by comparison to previously published datasets. For better characterization of T cell subsets, the identified T cell clusters were further subclustered using the cluster_cells function with resolution of 1e-3. Differential expression between naïve and activated CD4+ T cells was determined using the fit_models function. The dotplot in Fig 6A was generated using Plotly.

A list of Hobit induced genes (Mackay et al., 2016) was kindly shared by Dr. Laura Mackay and Dr. Luke Gandolfo. We then performed the GSEA in Fig 6A using Broad Institute’s GSEA software (Subramanian et al., 2005), to check if Hobit induced genes (p<0.05, FDR<0.05) were significantly enriched in 6 dpi activated lung CD4 effectors compared to naïve CD4 T cells in the lung (differentially expressed genes ranked by % expression fold-change).

### Statistical analysis

Unpaired, two-tailed, Students t-test was used to assess statistical significance between the means of two groups, with P < 0.05 considered significant. Analysis was done using Prism (Graphpad) software. Error bars in the figures represent the standard error of the mean. Significance in the figures are indicated as * *P* < 0.05, ** *P* < 0.01 and *** *P* < 0.001. Expression levels of different markers analyzed by flow cytometry are shown as MFI (Median Fluorescence Intensity) or nMFI (normalized MFI). To correct for batch effects while pooling data from different experiments, we normalized MFI by dividing each data point within an experiment by the average MFI of the control group from that experiment. nMFI = MFI/(average MFI of the control group)

