## Supplemental Information for "Cytotoxic CD4 Development Requires CD4 Effectors to Concurrently Recognize Local Antigen and Encounter Infection-Induced IL-15"

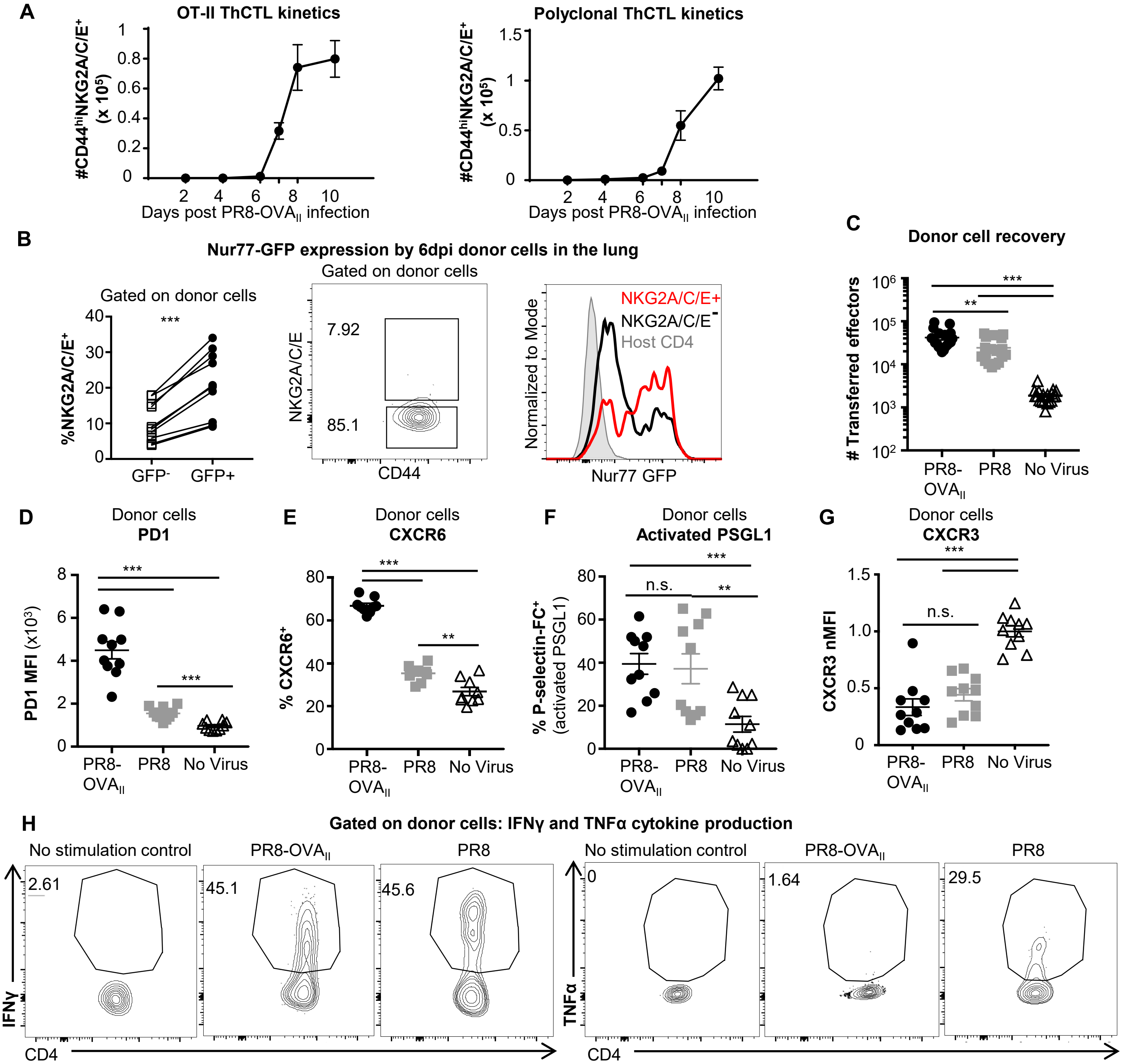


**Supplementary Figure 1 (Related to Fig. 1). Early ThCTL have higher levels of Nur77$^{GFP}$ expression corresponding to requirement for cognate Ag recognition. Kinetics of $T_{FH}$ generation during IAV infection. (A)** Naïve OT-II.Thy1.1$^{+}$ CD4 T cells were transferred into B6 mice and infected with PR8-OVA$_{II}$. Host polyclonal and donor OT-II lung ThCTL (NKG2A/C/E$^{+}$) generation was analyzed 2, 4, 6, 7, 8 and 10 dpi (n=7-8 pooled from 2 independent experiments). **(B)** Naïve OT-II.Nur77$^{GFP}$.Thy1.1$^{+}$ CD4 T cells were transferred into wild-type mice infected with PR8-OVA$_{II}$. At 6 dpi, donor cells in the lung were examined by flow cytometry. The percentage of GFP$^{+}$ and GFP$^{-}$ donor cells expressing ThCTL marker - NKG2A/C/E$^{+}$ (left). Representative FACS plot of NKG2A/C/E expression (middle). Representative histogram of GFP expression by NKG2A/C/E$^{+}$ donor cells, NKG2A/C/E$^{-}$ donor cells, or host CD4 T cells (right). **(C)** Experiment performed as in Figure 1A for (C-G). Donor cells in the lung were analyzed 2 dpt (8 dpi) by flow cytometry. Numbers of donor cells (n=19 per group pooled from 4 independent experiments). **(D)** PD1 MFI of donor cells. **(E)** Percentage of donor cells expressing CXCR6. **(F)** Percentage of donor cells that bind P-selectin. **(G)** Normalized CXCR3 MFI of donor cells. (D-G, n=9-10 per group pooled from 2 independent experiments). **(H)** Representative FACS plots of intracellular cytokine staining corresponding to Figure 1F-G. IFNγ and TNFα expression gated on donor lung cells that were restimulated with anti-CD3/anti-CD28 ex-vivo or unstimulated, corresponding to Figure 1F-G (n=7 per group pooled from 2 independent experiments). Error bars represent s.e.m. Statistical significance determined by two-tailed, unpaired Student's t-test (* $P < 0.05$, ** $P < 0.01$ and *** $P < 0.001$).

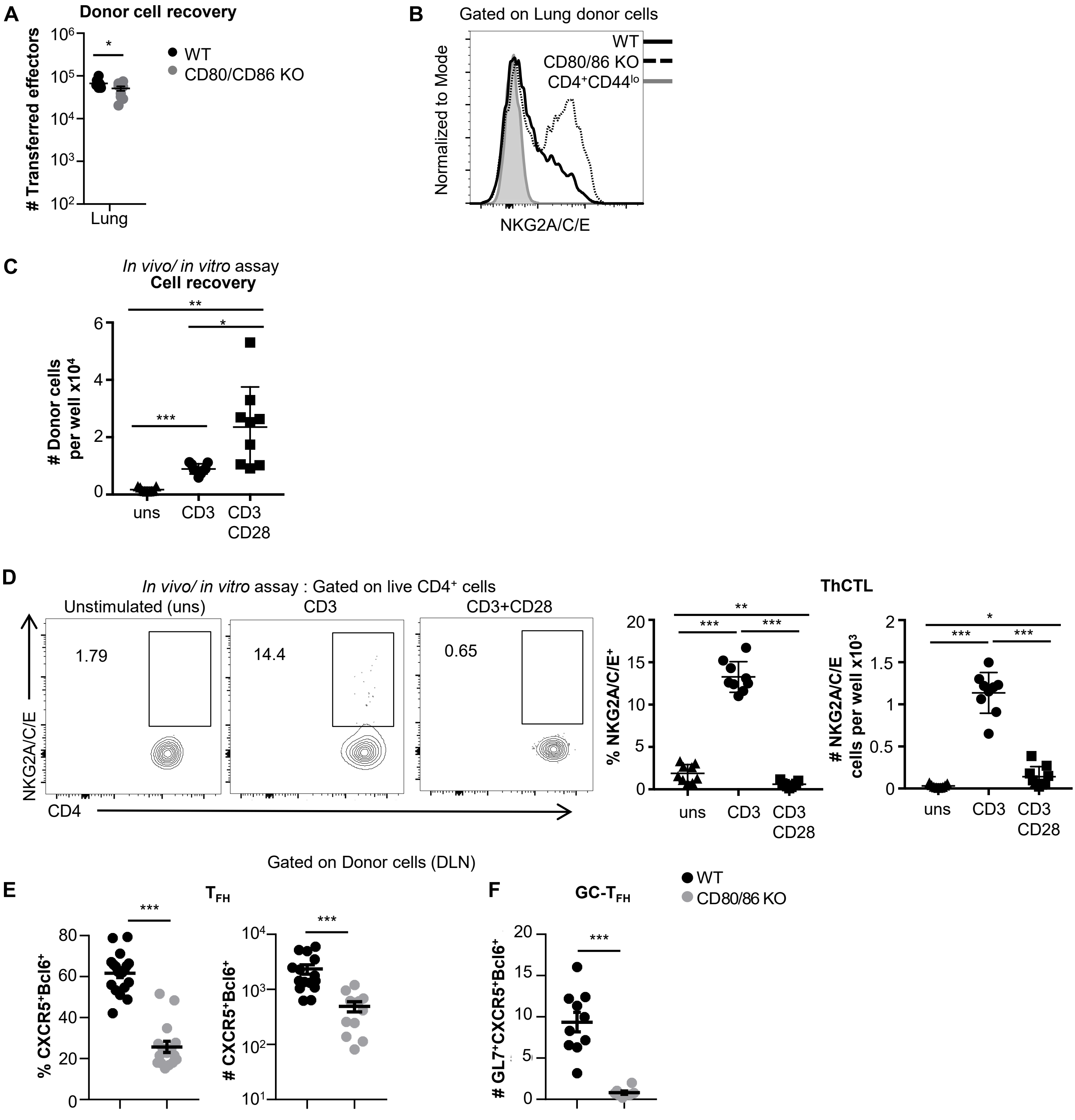


**Supplementary Figure 2 (Related to Fig. 2). *In vitro* TCR stimulation is sufficient to drive ThCTL differentiation from 6d *in vivo* generated effectors, while CD28 co-stimulation is suppressive. DLN $T_{FH}$ and GC-$T_{FH}$ generation from 6d effectors in the absence of CD28 co-stimulation *in vivo.*** **(A-B)** Experiment performed as in Figure 3A. **(A)** Number of donor cells in the lung (n=10 per group pooled from 2 independent experiments). **(B)** Representative histogram of NKG2A/C/E expression by donor cells isolated from lungs of WT mice or CD80/CD86$^{-/-}$ mice and naïve CD4 T cells. (n=9 per group pooled from 2 independent experiments). **(C-D)** Experiment performed as in Fig 2C. **(C)** Number of donor cells recovered in each group. **(D)** Percentage and numbers of donor cells expressing NKG2A/C/E after 2 days *in vitro* in the indicated conditions. (Cells were cultured in triplicate per condition and results are pooled from 3 independent experiments). **(E-F)** Experiment was performed as in Figure 2A. **(E)** Percentage and numbers of $T_{FH}$ generated from donor cells in the DLN **(F)** Number of donor DLN GC-$T_{FH}$. (n= 10-19 per group pooled from 2-4 independent experiments). Error bars represent s.e.m. Statistical significance determined by two-tailed, unpaired Student's t-test (* $P < 0.05$, ** $P < 0.01$ and *** $P < 0.001$).

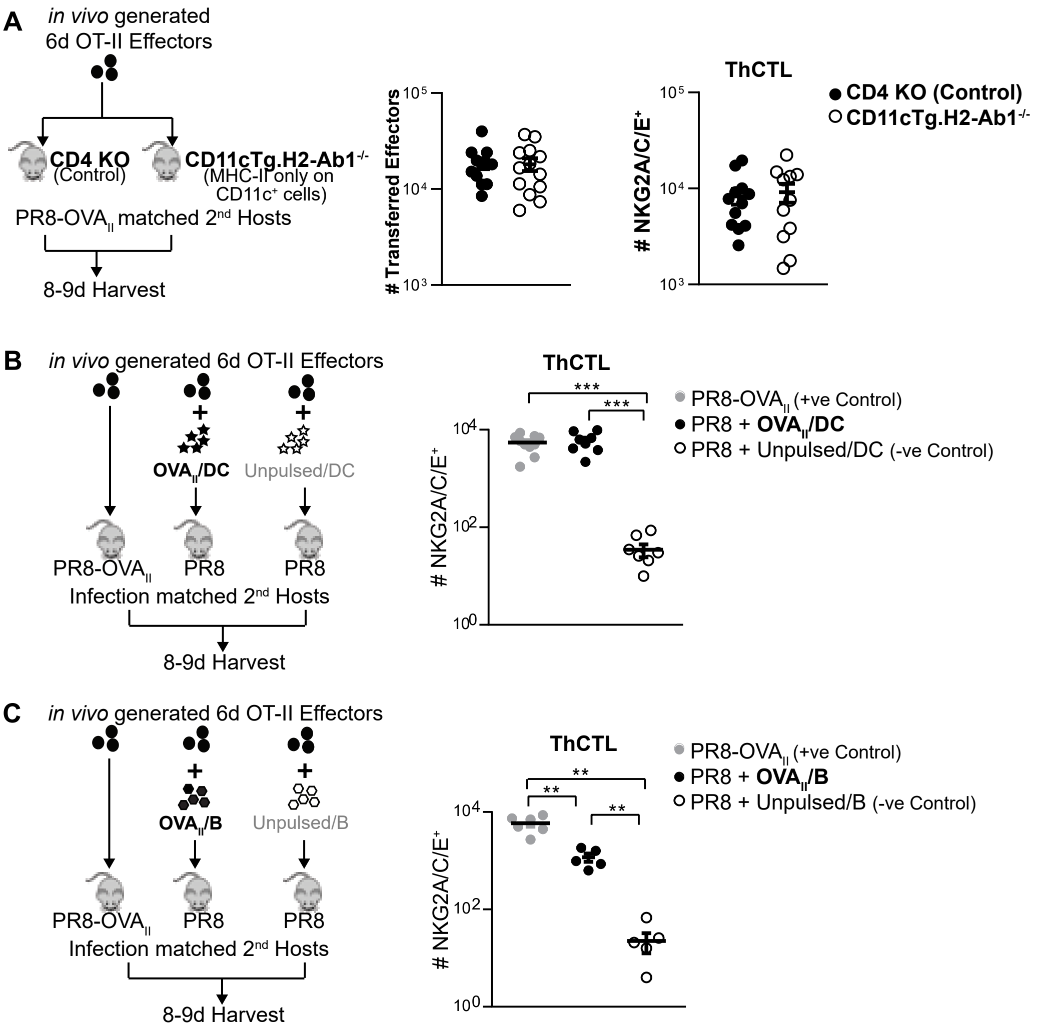


**Supplementary Figure 3 (Related to Fig. 3). DCs and B cells are able to support ThCTL generation from 6d effectors. (A)** *In vivo* generated 6d OT-II.Thy1.1$^+$ effectors were transferred into PR8-OVA$_{II}$ infection-matched CD11cTg.H2-Ab1$^{-/-}$ mice where MHC-II is restricted to CD11c$^+$ cells or into CD4 KO control mice. Total number of donor cells in the lung and ThCTL generation was analyzed at 8-9 dpi (n=7-11 per group pooled, 2-3 independent experiments). **(B-C)** *In vivo* generated 6d OT-II.Thy1.1$^+$ effectors were transferred into PR8-OVA$_{II}$ infection-matched as positive controls or into PR8 infection-matched into WT mice **(B)** with cognate Ag supplied via OVA$_{II}$ pulsed BMDC vs unpulsed BMDC controls (n=8-10 per group pooled, 3 independent experiments). **(C)** with cognate Ag supplied via OVA$_{II}$ pulsed B cells vs unpulsed B cell controls (n=5-6 per group pooled 2 independent experiments). The APC were transferred both intranasally and intravenously. Error bars represent s.e.m. Statistical significance determined by two-tailed, unpaired Student's t-test (* P<0.05, ** P<0.01 and *** P<0.001).

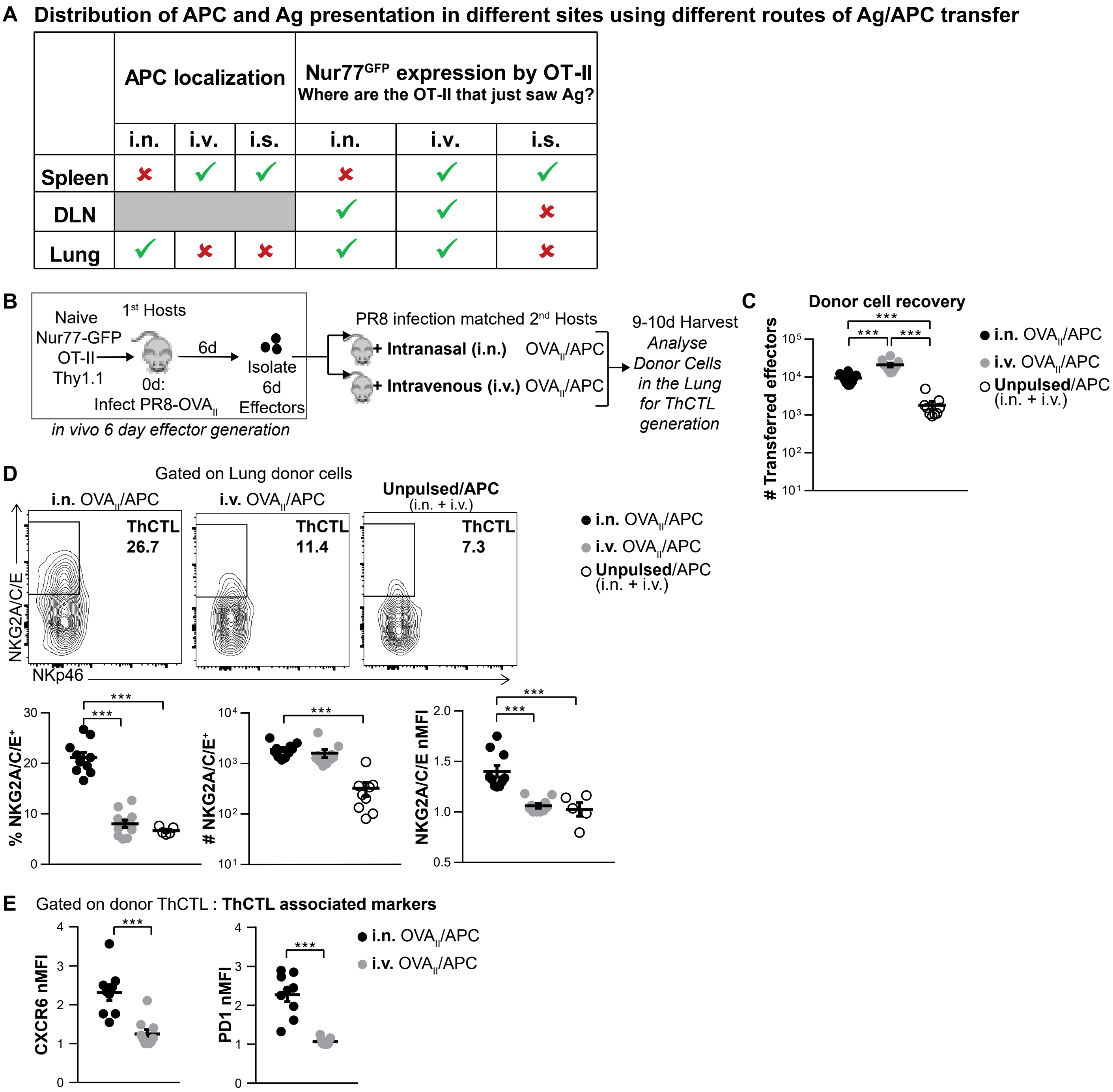


**Supplementary Figure 4 (Related to Fig. 4). Intravenous Ag delivery during the effector phase does not optimally support lung ThCTL generation from effectors. (A)** Distribution of APC and Ag presentation in different sites using intranasal (i.n.), intravenous (i.v.) or intrasplenic (i.s.) transfer of APC. Data summarized from Devarajan et al., 2022. $OVA_{II}$ peptide pulsed B-cells ($GFP^+$ or $CD45.1^+$) were used as APC and transferred into PR8 infection-matched hosts 6 dpi either intranasally (i.n.), intrasplenically (i.s.) or intravenously (i.v.). *In vivo* generated 6d OT-II.$Nur77^{GFP}$.$Thy1.1^+$ effectors were transferred i.v. Mice were harvested 14-16hr post-transfer (pt) and donor cells were analyzed by flow cytometry for donor $Nur77^{GFP}$ expression or for number of transferred APC (i.n. vs i.s.). **(B)** Experimental design: $OVA_{II}$ peptide pulsed B-cells were used as APC and transferred into PR8 infection-matched hosts 6 dpi either intranasally (i.n.) or intravenously (i.v.). Unpulsed APC were transferred both i.n. and i.v. as negative controls. *In vivo* generated 6d OT-II.$Nur77^{GFP}$.$Thy1.1^+$ effectors were transferred i.v. Mice harvested 3-4 dpt and ThCTL generation from donor cells in the lung was analyzed. **(C)** Number of donor effectors recovered with i.n. vs i.v. APC transfer. **(D)** Percentage and numbers of donor lung ThCTL formation with i.n. vs i.v. APC. nMFI of NKG2A/C/E expression of donor ThCTL generated. **(E)** Expression of ThCTL associated markers (CXCR6 and PD1) by donor ThCTL generated. (n=5-11 per group pooled, 3 independent experiments) Error bars represent s.e.m. Statistical significance determined by two-tailed, unpaired Student's t-test (* $P < 0.05$, ** $P < 0.01$ and *** $P < 0.001$).

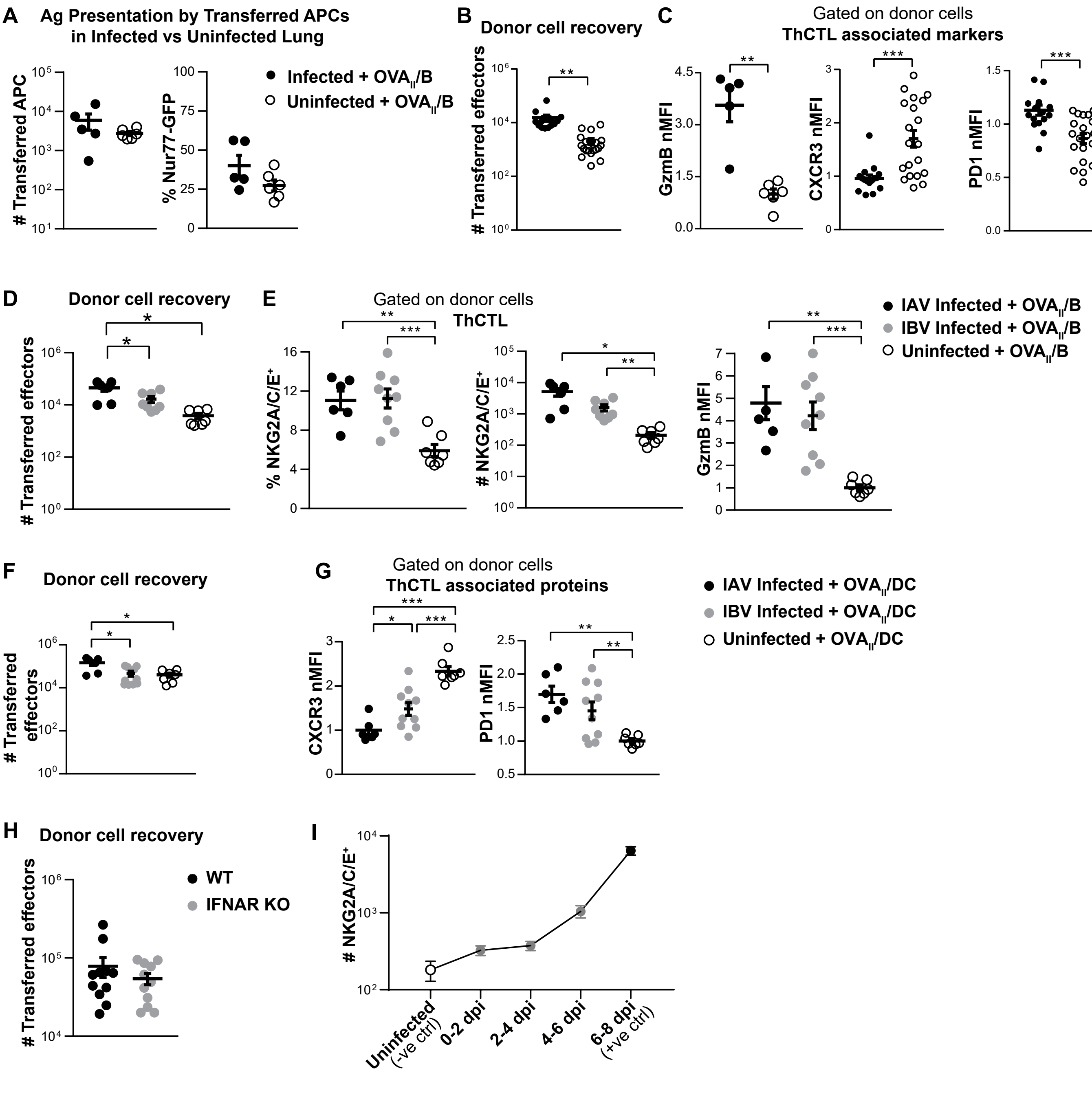


**Supplementary Figure 5 (Related to Fig. 5). Lung ThCTL generation is stunted in the absence of signals from infection after 6d (A)** CD45.1 $OVA_{II}$/B were transferred i.n. and i.v. into infection-matched hosts 6 dpi or into uninfected mice with in vivo generated 6d OT-II.Nur77GFP.Thy1.1$^{+}$ effectors. Numbers of transferred APC and Nur77$^{GFP}$ expression was assessed 14-16 hr pt. (n=5-6 per group pooled from 2 independent experiments). **(B-C)** Related to Fig 5B **(B)** Number of donor effectors recovered in the lung **(C)** Normalized MFI of ThCTL associated markers expressed – GzmB, CXCR3 and PD1 - by total donor cells in the lung. (n=16-20 per group pooled from 5 independent experiments). **(D-E)** 6d effectors + $OVA_{II}$/B were transferred into IAV or IBV infection-matched, or uninfected hosts. 4 dpt lungs were analyzed by FACS for **(D)** Number of donor effectors recovered in the lung **(E)** Donor ThCTL generation and GzmB expression by total lung donor cells. **(F-G)** Related to Fig 5C-D. **(F)** Number of donor effectors recovered in the lung. **(G)** ThCTL associated marker expression by donor lung effectors. (D-G, n=6-10 per group pooled from 2 independent experiments). **(H)** Related to Fig 5E. Lung donor cell recovery. (n=11-17 per group pooled from 3-4 independent experiments). **(I)** Related to Fig 5F. Number of ThCTL generated from donor effectors. (n=5-7 per group pooled from 2 independent experiments). Error bars represent s.e.m. Statistical significance determined by two-tailed, unpaired Student's t-test (* $P < 0.05$, ** $P < 0.01$ and *** $P < 0.001$).

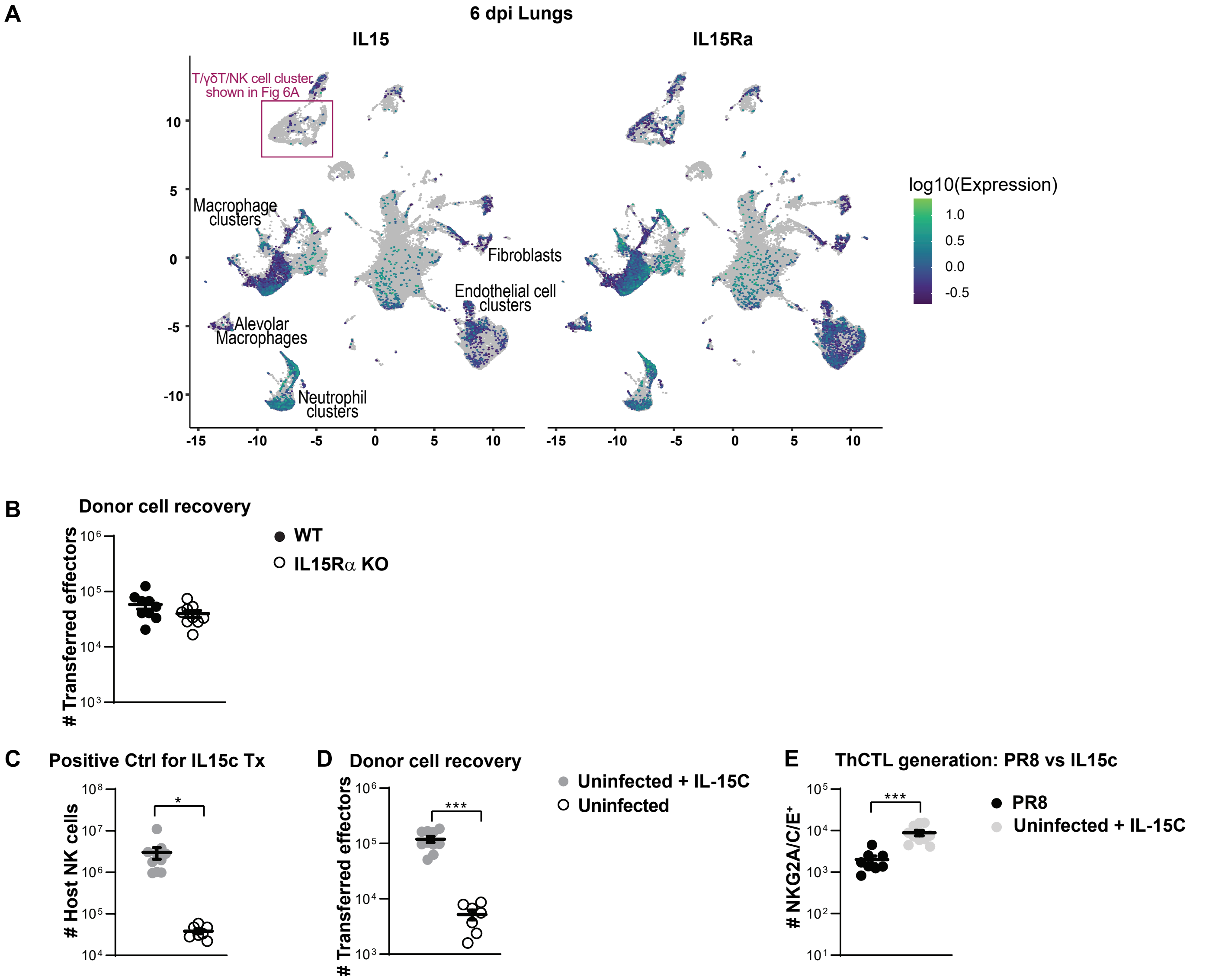


**Supplementary Figure 6 (Related to Fig. 6). Impact of IL-15. (A)** Influenza infected lungs were analyzed by scRNAseq at 6 dpi. Expression of IL15 and IL15Ra was analyzed and is shown in the UMAP plot. **(B)** Experiment was done as in Fig 6C. Total lung donor cell recovery is shown 2 dpt. (n=9-10 per group pooled, 2 independent experiments). **(C-D)** Experiment was done as in Fig 6E. **(C)** Number of host NK cells in the lung were analyzed as a positive control for IL-15c Tx after harvest 3 dpt. **(D)** Total donor cell recovery in the lung is shown 3 dpt. (n=8-10 per group pooled, 2 independent experiments). **(E)** Experiment was done as in Fig 6C and in addition *in vivo* generated 6d OT-II.Thy1.1$^{+}$ effectors were also transferred along with $OVA_{II}$/B cells into PR8 infection-matched hosts. The number of donor ThCTL generated 3 dpt was compared between IL-15c Tx hosts and PR8 infection-matched hosts. (n=8-10 per group pooled, 2 independent experiments). Error bars represent s.e.m. Statistical significance determined by two-tailed, unpaired Student's t-test (* $P < 0.05$, ** $P < 0.01$ and *** $P < 0.001$).
